# Circulation of enterotoxigenic Escherichia coli (ETEC) isolates expressing CS23 from the environment to clinical settings

**DOI:** 10.1101/2023.02.08.527665

**Authors:** Carla Calderon Toledo, Astrid von Mentzer, Jorge Agramont, Kaisa Thorell, Yingshun Zhou, Miklós Szabó, Patricia Colque, Inger Kuhn, Sergio Gutiérrez-Cortez, Enrique Joffré

## Abstract

Enterotoxigenic *Escherichia coli* (ETEC) is one of the leading causes of infant diarrhea in low- and middle-income countries (LMICs). Diarrheal pathogens are transmitted through environmental reservoirs; however, the bacterial clones that spread across the human-environment interphases remind unexplored. We aimed to determine the relationship and clonal dissemination of ETEC between children with diarrhea (> 5 years of age) and polluted water samples from local river in La Paz, Bolivia. Our study used whole genome sequencing and phenotypic fingerprinting system (PhenePlates) to analyze ETEC strains. We showed that ST218 and ST410 LT+STh CS23 ETEC were found with high frequency in both samples. The CS23 ETEC isolates were found within several STs, *E. coli* phylogroups A, B1, C, and D, and across ETEC lineages. Our comparative genomic analysis and PhenePlate screening of globally distributed clinical ETEC strains suggested that virulent CS23 plasmids acquisition occurs independently of the bacterial chromosomal background. Environmental strains were more often multidrug-resistant (MDR) than clinical isolates and harbored the class 1 integron-integrase gene *intI1* next to the MDR cassettes. Retrospective analysis of antibiotic resistance in ETEC revealed a high frequency of MDR in clinical isolates. The LT+STh CS23 ETEC isolates showed an increased biofilm ability at environmental temperature, equal cytotoxicity, and significantly lower adherence to human epithelial cells compared to ETEC expressing other CFs. Together, our findings suggest that CS23 is more prevalent in ETEC than previously estimated, and the Choqueyapu River is a reservoir for LT+STh CS23 ETEC containing strains capable of causing diarrheal cases in children.

**Importance:** The importance of clean water cannot be overstated. It is a vital resource for maintaining health and well-being. Unfortunately, water sources contaminated with fecal discharges from animal and human origin due to a lack of wastewater management poses a significant risk to communities, as they can become a means of transmission pathogenic bacteria like enterotoxigenic *E. coli* (ETEC). ETEC is frequently found in polluted water in countries with a high prevalence of diarrheal diseases, such as Bolivia. This study provides novel insights into the circulation of ETEC between diarrheal cases and polluted water sources in areas with high rates of diarrheal disease. The findings highlight the Choqueyapu River as a potential reservoir for emerging pathogens carrying antibiotic-resistance genes, making it a crucial area for monitoring and intervention. Furthermore, the results demonstrate the feasibility of a low-cost, high-throughput method for tracking bacterial pathogens in low- and middle-income countries, making it a valuable tool for One Health monitoring efforts.

## Introduction

Enterotoxigenic *E. coli* (ETEC) is a food- and waterborne pathogen and one of the leading causes of moderate to severe diarrhea in children, particularly in low- and middle-income countries (LMICs) and adults traveling to endemic regions (1, 2). The Global Burden of Disease study included ETEC among the top ten diarrheal-causing agents in the world, accounting for more than 50,000 annual deaths and 223 million cases per year (3). ETEC infection causes varying symptoms, from mild diarrhea to a severe cholera-like disease, by colonizing the small intestine using colonization factors (CFs) and the secretion of heat-labile toxin (LT) and/or heat-stable toxins (STh and STp). Over 30 different ETEC CFs have been described and shown to be associated with human infections (4–6), being the most common CFA/I, CS1-7, CS14, CS17 and CS21 (5). However, toxin-positive but CF-negative ETECs are frequently detected in epidemiological studies (7, 8).

Transmission of ETEC occurs via the fecal-oral route by consuming contaminated food and water (9). Studies in Bolivia have shown that not only is ETEC one of the most prevalent diarrheagenic *E. coli* pathotypes that infect children under 5 years of age (10), but high levels of ETEC bacteria are also found in agricultural soils, farm crops, and river waters (11, 12). Since ETEC can survive for a long time in water, environmental transmission is a risk (1, 13, 14). It is unknown to what extent clinical cases have a water or food origin in La Paz, but it has been suggested that the environmental origin is highly plausible (10). La Paz, with approximately 1 million inhabitants, does not have a wastewater treatment plant, and wastewater is discharged directly into the Choqueyapu River. Downstream of the Choqueyapu River, many communities use river water for crop irrigation, whose products are mainly commercialized in markets in the city of La Paz (11, 12). In a recent global study of water pollution, some rivers in the La Paz Basin were reported to be among the most polluted rivers, with high levels of pharmaceutical residues (15). These reports highlight the need to carry out One Health strategy studies that help explain the largely unknown interactions between humans and the environment in the emergence and transmission of diarrheal diseases.

In the present study, we characterize ETEC isolates from diarrheal cases of hospitalized children and from water samples collected on the Choqueyapu River in La Paz, Bolivia. Whole genome sequencing was used to determine the relationship and clonal dissemination of ETEC between clinical and environmental samples. We also compared the genomic results of ETEC diversity and the relationship of bacterial strains from different sources using a low-cost phenotypic fingerprinting system (PhenePlate or PhP method). We found that a surprisingly high number of ETEC isolates harbored a less common CF, CS23, which is usually undetected due to the lack of established detection methods for this CF. Therefore, CS23-positive isolates are grouped into CF-negative ETEC in most studies but may be more prevalent in the environment and clinical settings than previously anticipated.

## Results

### Whole genome analysis of clinical and environmental strains reveals that CS23 circulates in both settings

Clinical and environmental isolates were collected in 2013-2014 and 2014, respectively, to determine the spread of ETEC between the two interfaces and characterize the ETEC isolates using whole genome sequencing (Table 1 and Table S1). In total, 30 ETEC strains with positive PCR results for ETEC toxins were included in the study, of which 11 isolates were isolated from hospitalized children (13.2 ± 8.2 months) with acute diarrheal disease. During the same sampling period, 19 isolates were obtained by filtering shallow water at six different sites along the Choqueyapu River, where its waters are used for agricultural irrigation (Table S1). All isolates positive for toxins in PCR were whole genome sequenced, and genes and gene clusters encoding toxins (LT and/or STh or STp) and colonization factors were identified and extracted using an open-source database (https://github.com/avonm/ETEC_vir_db). Eleven combinations of toxins / CFs were identified; Clinical isolates reported a greater diversity of virulence factors that accounted for eight virulence profiles, while environmental strains displayed four different profiles (Table 1). We found that LT + STh CS23 was the most prevalent virulence profile among environmental isolates (15/19), and the same profile was found in three (3/11) clinical isolates. Four isolates were found to harbor a cluster of genes that encode a putative chaperone-usher assembled pili, where three of the isolates corresponded to clinical samples. Comparison of the CS23 major subunit (AalE) extracted from the isolates with the CS23-variant revealed five major AalE types (AalE-1-5) (Fig. S1 and Table S1). The CS23 variant AalE-2 (ET13, ET18 and M5KL (1)) and AalE-5 (ET11, ET15, ET21, ET22, ET26 and 149 B7(1)) were found among clinical and environmental samples. The other CS23 variants, AalE-1 and AalE-4, were present exclusively in a few environmental isolates, while AaE-3 was found in the clinical isolate MALC (1) (Fig. S1 and Table S1). Regarding the other nine clinical isolates, they had virulence profiles commonly reported in the literature (CS1+CS2+CS21, CS1+CS3+CS21, CS6+CS8, and CS21-only) (16, 17). Additional characterization of non-classical ETEC virulence factors identified two serine proteases encoded by *eatA* and the two-partner apparatus *etpBAC* of the EtpA fimbria (Table 1).

**Table 1.** Phenotypic and genomic characteristics of ETEC isolates collected in Bolivia.

| Isolate ID | Source | Type | Toxin Profile | CF profile | Non-classical VF | MLST (ST) | Phylo-group | ETEC lineages |
| --- | --- | --- | --- | --- | --- | --- | --- | --- |
| ET5 | Water | River | LT*+STh | CS23<sup>1</sup> |  | 69 | D | ND |
| ET6 | Water | River | LT | Putative CU-pili |  | 155 | B1 | 19 |
| ET7 | Water | River | LT+STh | CS23 |  | 58 | B1 | 19 |
| ET8 | Water | River | LT+STh | CS23 |  | 949 | B1 | 19 |
| ET9 | Water | River | LT+STh | CS23 |  | 410 | C | 16 |
| ET10 | Water | River | LT+STh | CS23 |  | 58 | B1 | 19 |
| ET11 | Water | River | LT+STh | CS23 |  | 218 | A | 12 |
| ET12 | Water | River | LT+STh | CS23<sup>1</sup> |  | 949 | B1 | 12 |
| ET13 | Water | River | LT+STh | CS23 |  | 410 | C | 16 |
| ET14 | Water | River | STh | CS23 |  | 10 | A | 15 |
| ET15 | Water | River | LT+STh | CS23 |  | 949 | B1 | 19 |
| ET17 | Water | River | LT+STh | CS23<sup>1</sup> |  | 10 | A | 15 |
| ET18 | Water | River | LT+STh | CS23 |  | 410 | C | 16 |
| ET19 | Water | River | LT+STh | CS23 |  | 10 | A | 15 |
| ET21 | Water | River | LT*+STh | CS23 |  | 218 | A | 12 |
| ET22 | Water | River | LT*+STh | CS23 |  | 949 | B1 | 19 |
| ET23 | Water | River | LT | <sup>2</sup>CS6+CS8 | *eatA* | 3931 | A | 4 |
| ET25 | Water | River | LT | <sup>2</sup>CS6+CS8 | *eatA* | 3931 | A | 4 |
| ET26 | Water | River | LT+STh | CS23<sup>1</sup> |  | 949 | B1 | 19 |
| 51 LAE | Clinical | AD | LT | Putative CU-pili | *etpBC* * | 10 | A | 15 |
| 78 CRE+ | Clinical | AD | LT+STh | <sup>2</sup>CS2+CS3<sup>3</sup>+CS21 | *eatA*<br>*etpBC* * | 4 | A | 2 |
| 149 B7 | Clinical | AD | LT+STh | CS23 |  | 218 | A | 12 |
| 169 LAF | Clinical | AD | LT | CS6 |  | 10 | A | 15 |
| 250 B7 (1) | Clinical | AD | LT<sup>§</sup> | Putative CU-pili |  | 155 | B1 | 19 |
| 284 B7 (1) | Clinical | AD | STh | CS21 |  | 48 | A | 4 |
| 570 LAE+ | Clinical | AD | LT | CS1+CS3+CS21 |  | 4 | A | 1 |
| 619 B8 (1) | Clinical | AD | STh | CS21 |  | 48 | A | 4 |
| 724 B9 (1) | Clinical | AD | LT | CS13 + putative<br>CU-pili | *eatA*<br>*etpBC* * | 46 | A | 22 |
| M5 KL (1) | Clinical | AD | LT+STh | CS23 |  | 410 | C | 16 |
| MALC (1) | Clinical | AD | LT+STh | CS23 |  | Novel | A | 15 |

AD: acute diarrhea.
VF: virulence factors
(\*) LT gene duplicated; § (1) interrupted operon (could be due to sequencing issues); (2) *cfaD* gene present; (3) missing *cstB* gene from the CS3 operon.
\* Disrupted contigs

### Circulation of CS23 ETEC isolates with the same ST group between clinical and environmental settings

To assess the heterogenicity of ETEC strains and their genetic relationship, we performed *in silico* multilocus sequence typing (MLST) and *E. coli* phylogroup typing (Table 1). First, the MLST results showed that our 30 ETEC strains belonged to 11 different sequence types (STs), of which three were found only in clinical isolates (ST4, ST48, and ST46) and four only in environmental isolates (ST58, ST69, ST949, and ST3931). Furthermore, four STs were found in clinical and environmental samples, including the presumed high-risk clone of *E. coli* ST410 and ST218, ST155, and ST10. Based on virulence profiles, LT + STh CS23 strains belonged to six ST groups (ST58, ST69, ST218, ST410, ST949, and ST10). Therefore, CS23 was not associated with a specific ST type or clonal group, indicating that CS23 is located on a mobile plasmid.

The 30 ETEC strains analyzed were highly diverse, since they fall into four phylogroups of *E. coli* (A, B1, C, and D) (Table 1). Phylogroup A was primarily associated with clinical isolates, while phylogroups A, B1, and C were represented among environmental isolates. The ET5 isolate was the only strain that belonged to phylogroup D. Phylogroup C included clinical and environmental strains, and all were ST410. CS23-positive ETECs were not restricted to a particular phylogroup. On the other hand, strains expressing CS1-CS3, CS6, CS8, CS13, or CS21 were limited to the A phylogroup that supports our previous findings (18).

Phylogenetic analysis of the 30 ETEC isolates showed a consistent clustering pattern with MLST groups and demonstrated specific clusters of environmental and clinical ETEC with a high degree of genetic relatedness (Figure 1a). The phylogenetic tree topology grouped the ETEC genomes into clusters A and B. Cluster A consisted of 11 environmental and two clinical genomes, grouped into subgroups of ST949, ST155, ST58 and ST410. The ST155 and ST410 groups within this cluster included both environmental water and diarrheal isolates. Cluster B included 9 of 11 clinical isolates and seven environmental isolates. The subgroup of ST218 genomes included isolates from mixed sources. ST10 isolates were both clinical and environmental, but they were not grouped together according to the phylogenetic analysis. Instead, subclusters of solely clinical and environmental genomes were formed. The environmental ST3931 isolates were closely related to clusters of clinical ST4 and ST48 isolates (Figure 1a). Based on the virulence profile, the LT+STh CS23 genomes were spread across the A and B clusters. Interestingly, the ET5 LT+STh CS23 strain was the most genetically distant isolate of all. Overall, our data confirm the presence of two LT+STh CS23 ETEC clones, ST410 and ST218, in the waters of the Choqueyapu River that cause diarrheal cases during the time of sampling. Furthermore, expected high genetic diversity among clinical isolates and close relatedness between clinical and environmental isolates were observed.

**Figure 1.**
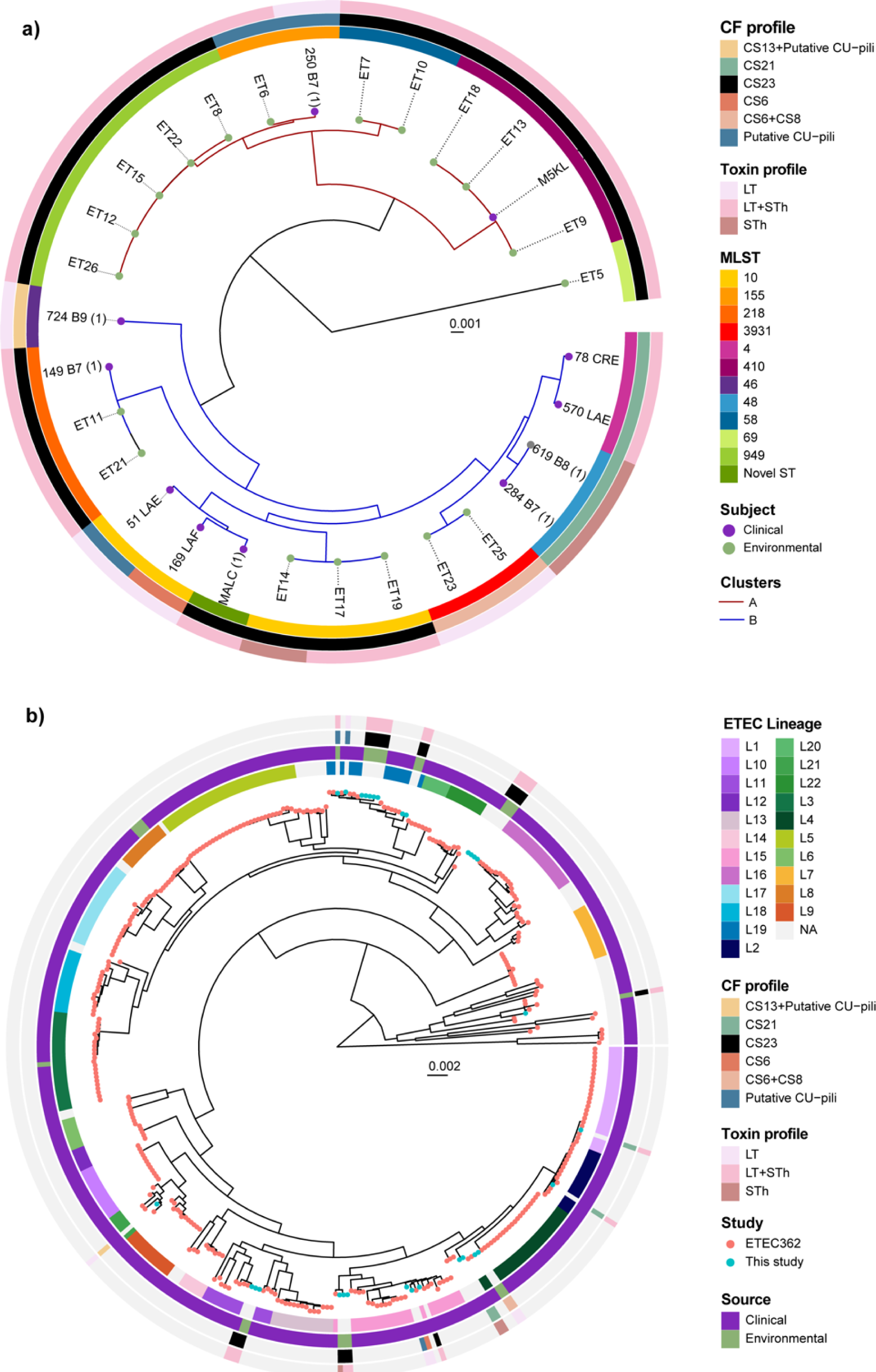
Whole-genome phylogenetic analysis of ETEC. Maximum likelihood midpoint-rooted phylogenetic trees based on SNP differences in **(a)** 30 ETEC genomes collected in Bolivia and **(b)** in context with 362 selected ETEC genomes with representative virulence profiles isolated from indigenous patients and travelers between 1980 and 2011 from endemic countries (Asia, Africa and North, Central and South America). The tip of the branches is color-coded according to the sample’s origin (clinical/environmental in this study and the 362 reference ETEC genomes). The colored rings represent the respective MLST and CF profiles, and the lineages L1-L21 are indicated. The scale bar represents the substitutions per variable site.

To identify whether Bolivian strains belonged to previously described ETEC lineages, we built a phylogenetic tree in which the reference genomes of 362 clinical ETEC strains were included (Figure 1b). The collection of 362 ETEC genomes was obtained from children and adults with diarrheal disease from endemic countries (Asia, Africa and Central and South America) for 30 years (1980 to 2011) (18). The results demonstrated that our new collection of Bolivian ETEC strains belonged to eight previously described ETEC lineages (L1, L2, L4, L12, L15, L16, L19 and L22) (18). Surprisingly, all environmental isolates were genetically close to clinical ETEC strains distributed worldwide. Only six of the Bolivian strains of this study (570_LAE, 78_CRE, 284_B7, 619_B8, ET23, and ET25) belonged to the major lineages of ETEC (L1 - L5), including the most prevalent virulence profiles. Their virulence profiles were consistent with the virulence profile associated with each lineage in the von Mentzer study (18). For example, L4, which commonly encompasses ETEC strains expressing CS6, CS6+CS8, and CS21, also clustered with the STh CS21 and LT CS6+CS8 ETEC isolates from this study. Interestingly, ETEC strains expressing CS23, CS6, CS13, and CF-negative fell into ETEC lineages (L12, L15, L16, L18 and L22) where most of the strains are CF-negative, *i.e.,* do not express any of the known ETEC CFs. The number of negative CF ETECs is partly due to the lack of standardized detection methods for many CFs, such as CS23, but may also indicate the presence of new CFs to be discovered. Similarly to the MLST results, the lineage identification of the LT + STh CS23 genomes indicated that these strains were distributed across four ETEC lineages (L12, L15, L16 and L19). A representative ETEC LT + STh CS23 isolated from a diarrheal case from the von Mentzer reference collection was found in L11. These results showed that environmental isolates share high levels of genetic similarity with clinical isolates associated with acute diarrhea cases around the world. Furthermore, we showed that LT + STh CS23 ETEC isolates can be found in highly heterogeneous genetic backgrounds and, in light of the ubiquitous findings of CS23 ETEC in this study, this CF should be considered in future studies of ETEC diarrhea.

### Antimicrobial susceptibility profiles, resistance genes, and incompatibility groups

To test for the presence of antibiotic resistance, an antimicrobial susceptibility test (AST) of the 30 ETEC isolates was performed with 16 antimicrobials belonging to nine classes of antibiotics. The results showed phenotypic resistance to 11 of 16 antimicrobials tested (Figure 2). In general, 90% of clinical isolates and 95% of environmental isolates were resistant to at least one class of drugs. Neither of the isolates detected resistance to β-lactams (cefotaxime, cefpodoxime, meropenem, imipenem) nor resistance to gentamicin. Multidrug resistance (MDR, resistance to 3 or more classes of antibiotics) was more prevalent in clinical isolates (63%) than in environmental isolates (52%). Resistance to streptomycin (STM) was the only antibiotic resistance significantly more frequent in clinical isolates than in environmental isolates (73% vs. 26%, Fisher p-value = 0.0119) (Table S2). Resistance to trimethoprim-sulfamethoxazole (TMP–SMX) was detected twice more frequently in clinical isolates (64%) than in environmental isolates (32%). Among environmental isolates, ampicillin resistance was the most frequently identified (52%), and comparable frequencies were also detected in clinical isolates (45%). Although resistance to fluoroquinolones such as ciprofloxacin - a widely used antibiotic for the treatment of acute diarrhea in Bolivia (19) - was found to be low in both isolate groups, resistance to nalidixic acid was found in 45% of the isolates. Based on the classification of ST clonal groups, ST groups with the highest average (μ) of phenotypic antibiotic resistance per strain are ST48 (μ = 4.5), ST3931 (μ = 4), ST949 (μ = 3.8), and ST10 (μ = 3.2). Interestingly, ST410, considered a high-risk MDR clone of *E. coli*, included non-MDR isolates with resistance to nalidixic acid and, in some cases, ciprofloxacin or amikacin. ETEC isolates sensitive to all antibiotics belonged to ST218, with the exception of ST21, which had resistance to amikacin (Figure 2).

**Figure 2.**
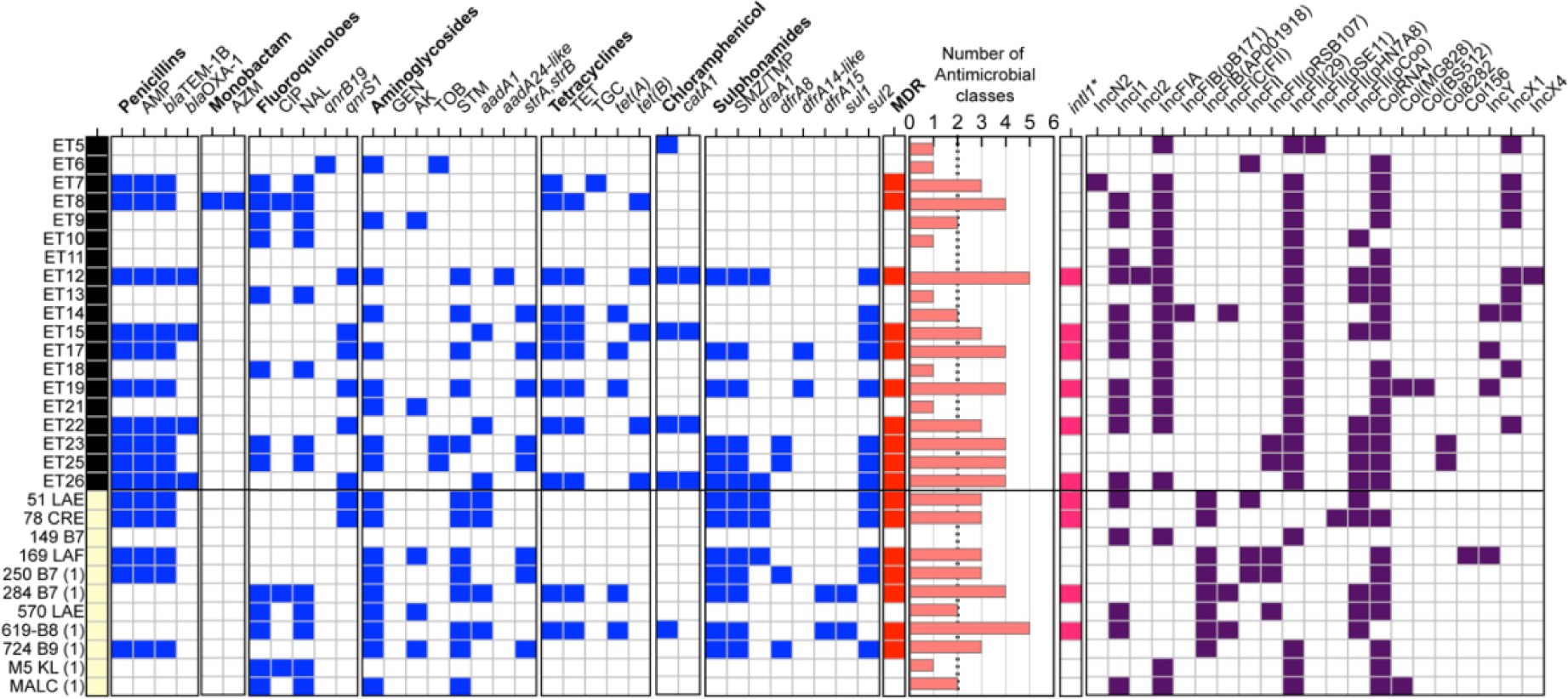
Phenotypic and genomic characterization of antibiotic resistance among ETEC isolates. Environmental (black) and clinical (yellow) samples were subjected to a phenotypic disc diffusion test (capital letters). Antimicrobial resistance genes were extracted from ETEC genomes (in italics). The blue boxes represent resistant/positive isolates, and the red boxes indicate multidrug resistance (≤3 classes of antibiotics). The histogram illustrates the number of antimicrobial classes in which the ETEC strains were phenotypically resistant. Genomic identification (≥90% nucleotide identity and ≥90% nucleotide identity) of the mobile element class 1 integron (*intI1*) (GenBank: AEH26333.1). Characterization of different Incompatibility Group (Inc) plasmids detected in bacterial genomes. AMP, ampicillin; AZM, azithromycin; CIP, ciprofloxacin; NAL, nalidixic acid; GEN, gentamycin, AK amikacin; TOB, tobramycin; STM, streptomycin; TET, tetracycline; TGC, tigecycline; CHL, chloramphenicol and TMP–SMX, trimethoprim–sulfamethoxazole.

Genomic screening of genes encoding antimicrobial resistance in the 30 ETEC genomes identified 17 antibiotic resistance genes (ARGs) that confer resistance to 5 different classes of antibiotics (Figure 2). Therefore, eight ARGs that confer resistance to aminoglycosides (*aadA1* and *strA* / *strB*), sulfonamides and trimethoprim (*sul2*, *drfA1* and *drfA8*), penicillin (*bla*_TEM-1b_), fluoroquinolones (*qnrS1*) and tetracyclines (*tetA*) were shared by environmental and clinical isolates. No statistical differences were found in the prevalence of ARGs between the two groups. The average number of ARGs between environmental and clinical isolates was similar (2.9 vs. 2.8), but a higher diversity of ARGs was observed in environmental strains, which exclusively contained the following ARG alleles: *aadA24*, *dfrA14-like*, *bla*_OXA-1_, *qnrB19, tetB,* and *catA1.* The sulfonamide trimethoprim resistance alleles *sul1* and *dfrA15* were only found in clinical isolates. No carbapenemases, rifampicin, macrolides, or colistin resistance genes were detected. According to our phenotypic antibiotic resistance profile, all ST218 (ET11, ET21, and 149 B7) and ST410 (ET9, ET13, ET18, and M5 KL) did not harbor any ARG. However, some were phenotypically resistant to fluoroquinolones, possibly due to the acquisition of a mutation in DNA gyrase and topoisomerase IV and not plasmid-borne resistance genes. As shown in Figure 2, the comparison of the AMR phenotype and genotype correlated to a large extent, specifically ampicillin-resistant strains (*bla*_TEM-1B_ *+ bla*_OXA-1_), tetracycline-resistant strains (*tetA* + *tetB*), trimethoprim–sulfamethoxazole-resistant strains (*draA1*, *draA8, dfrA15*, *sul1* and/or *sul2*) and chloramphenicol-resistant strains (*catA1*). Exceptions were observed in ET5 and 250 B7 isolates (1) that were negative for any chloramphenicol resistant gene.

As expected from the AST results, the β-lactam-resistant allele *bla*_TEM-1B_, which confers resistance to AMP, was the most frequent, followed by *sul2*, *strA* and *strB*, respectively. Another β-lactamase gene, *bla*_OXA-1,_ was found only in environmental isolates (ET12, ET15, ET22, and ET26). It is surprising to find that 13 ETEC isolates, consisting of six clinical strains and seven environmental, did not have any ARG. This includes all ST410 strains.

Further analysis of the genomic context of ARG of MDR clonal groups led to the localization of class 1 integron integrase (*intI1*) in the same vicinity as ARGs (Figure 2, Figure S2). Although ET12, ET26, and 51 LAE were positive for the *intI1* gene, the contigs were too small, so they contained only the class 1 integrase gene. The *intI1* gene is found in a variety of antibiotic resistant bacteria, both in the environment and in humans, and its presence can be used as an indicator of horizontal gene transfer, which occurs naturally in the environment or as the result of human activities such as antibiotic use and fecal pollution (20). Adjacent to *intI1*, two types of MDR cassettes were found in clinical and environmental strains. The first MDR cassette consisted of *sul1-qacEdelta1-addA1*(or *15*)*-dfraA1*(or 15)*-intI1* and was carried only by clinical isolates. As for the second type *addA1-bla*_oxa-1_-intl1 with *catA1* gene downstream of the cassette, only environmental genomes (ET15 and ET22) harbored this cassette. The ET17 and ET19 contigs were nearly identical (>98% identity) and contained the *dfraA14* allele next to the *intI1* gene (Figure S2). Blastn analysis of contigs carrying ARGs and *intl1* showed that most, but not all, contigs had high coverage and similarity to ETEC and *Enterobacterales* plasmids (*Shigella dysenteriae*, *Klebsiella pneumoniae* and *Escherichia coli*). For instance, blastn analysis of the contigs in 284 B7(1) and 619 B8 (1) showed high similarity with the L4_E1441_ ETEC plasmid 2 (pAvM_E1771_17 LR883013.1) characterized by carrying the *sul1* and *tetA* genes and the *Ing* operon encoding CF CS21 (21). Exceptionally, the ET22 MDR cassette was integrated into a 93,402-nucleotide long chromosomal contig (Figure S2).

As many virulence genes and ARGs are carried on mobile genetic elements, such as plasmids, we searched for plasmid replicon types (incompatibility groups) in ETEC isolates. Replicon typing identified a wide diversity of incompatibility groups such as IncF (IncFIA, IncFIB, IncFIC, and IncFII), IncN (IncN2), IncI (IncI1 and IncI2), Col-like (ColRNAI), IncY and IncX (IncX1 and IncX4). IncF was present in all ETEC strains, and the distribution of IncI, Col-like, and IncY was similar between clinical and environmental isolates, except IncX plasmids (ET5, ET7-9, ET12-14, ET18, and ET22) and IncN2 (ET7) that were only found in environmental strains. We did not find a particular association between the Inc plasmid group and ARGs; however, 100% of the LT + STh CS23 positive isolates harbored IncFIA+IncFII (*n*=29). Within the IncF plasmid, the pMLST types detected the following alleles: B18 (*n*=2), B53 (*n*=2), B45 (*n*=1), B52 (*n*=1), and A16:B45 (*n*=1). B18 variants were commonly detected in ST3931 isolates expressing LT CS6+CS8, while B53 was found in the 250 B7(1) clinical strains. The rest of the IncF plasmids had unknown STs.

### ETEC isolates expressing CS23 showed elevated biofilm formation at water temperatures

Biofilms are multicellular aggreges formed by surfaces on surfaces, used as a bacterial strategy to enhance persistence in aquatic environments and the host’s gastrointestinal tract (22). The colony morphotypes of ETEC isolates were tested at temperatures that mimic environmental conditions (20°C) and the mammalian gastrointestinal tract (GI) (37°C). Results from the Congo Red assay showed that 20°C induced the expression of the *rdar* morphotype (red, dry, and rough phenotype, expressing curli fimbriae and cellulose and indicative of elevated biofilm formation) in most of the ETEC isolates (14 environmental isolates and six clinical isolates) while at 37°C the *bdar* phenotype (brown, dry, and rough with curli expression) was commonly expressed by both clinical environmental isolates (Table 2 and Figure S3). A higher frequency of isolates with the saw morphotype (smooth and white expressing neither curli nor cellulose) was observed at 20°C than at 37°C. No association was found between the MLST groups and biofilm ability, although strains belonging to ST58, ST410, ST10, ST155, and ST949 were prompted to higher biofilm production than the ST218 and ST3931 strains. Of the 20 tested positive for rdar, 15 were found to be CS23 positive.

**Table 2.** Effect of temperature on the biofilm formation of ETEC isolates. Morphotype distribution in environmental and clinical isolates of ETEC. Morphotype: rdar (cellulose/curli fimbriae), pdar (cellulose/no fimbriae), bdar (no cellulose/curli fimbriae), and saw (no cellulose/no curli fimbriae). The ETEC strains were grown at 20°C or 37°C.

| Morphotype change | Environmental |  | Clinical |  |
| --- | --- | --- | --- | --- |
| 20°C → 37°C | n (%) | ST group (Isolate ID) | n (%) | ST group (Isolate ID) |
| *rdar*→*rdar* | 5 (26.3) | ST69 (ET5), ST410 (ET9), ST10 (ET14, ET17, ET19) | 3 (27.3) | ST155 (250 B7 (1)), ST218 (149 B7), ST410 (M5 KL (1)) |
| *rdar*→*bdar* | 6 (31.6) | ST155 (ET6), ST58 (ET7), ST949 (ET8, ET15, ET12), ST410 (ET13) | 2 (18.2) | ST10 (169 LAF), ST48 (619-B8 (1)) |
| *rdar*→*pdar* | 0 (0.0) |  | 1 (9.1) | ST48 (284 B7 (1)) |
| *rdar*→*saw* | 3 (15.8) | ST58 (ET10), ST949 (ET22, ET26) | 0 (0.0) |  |
| *bdar*→*rdar* | 1 (5.3) | ST419 (ET18) | 0 (0.0) |  |
| *bdar*→*pdar* | 0 (0.0) |  | 1 (9.1) | Novel ST (MALC (1)) |
| *bdar*→*saw* | 0 (0.0) |  | 1 (9.1) | ST10 (51 LAE) |
| *bdar*→*bdar* | 0 (0.0) |  | 2 (18.2) | ST4 (570 LAE), ST46 (724 B9 (1)) |
| *saw*→*saw* | 3 (15.8) | ST218 (ET21), ST3931 (ET23, ET25) | 1 (9.1) | ST4 (78 CRE) |
| *saw*→*bdar* | 1 (5.3) | ST218 (ET11) | 0 (0.0) |  |

### Clinical isolates exhibit higher adherence to mammalian cells, but similar cytotoxicity levels as environmental isolates

The adhesion of clinical and environmental ETEC isolates was evaluated in the Caco-2 epithelial human cell line. The results showed that clinical ETEC demonstrated a significant difference in clinical ETEC adhesion with an average of 38.5% compared to 2.1% from environmental isolates (*P* < 0.01) (Figure 3a). Given that CFs play a crucial role in ETEC binding to and colonizing the epithelial cells of the host, the isolates were sorted out according to their colonization factor profiles (positive and negative for CS23). For example, clinical isolates that express other CFs than CS23 (*i.e*., CS21, CS6, CS6+CS8, CS13, CS1, CS2, CS3, and putative CU-pili) had significantly higher binding ability compared to any other group (P < 0.05, Figure 3b). A large majority (11/19) ETEC isolates expressing CS23 from both groups were unable to attach to Caco-2 cells (< 1% of attached bacteria). This lack of adherence was also observed in a few isolates with the following CF profile: CS1+CS3+CS21 (284 B7 (1)), and CS6+CS8 (ET23 and ET25). These results showed the low ability of CS23 to mediate bacterial adherence to host epithelial cells and highlight the importance of other colonization factors.

**Figure 3.**
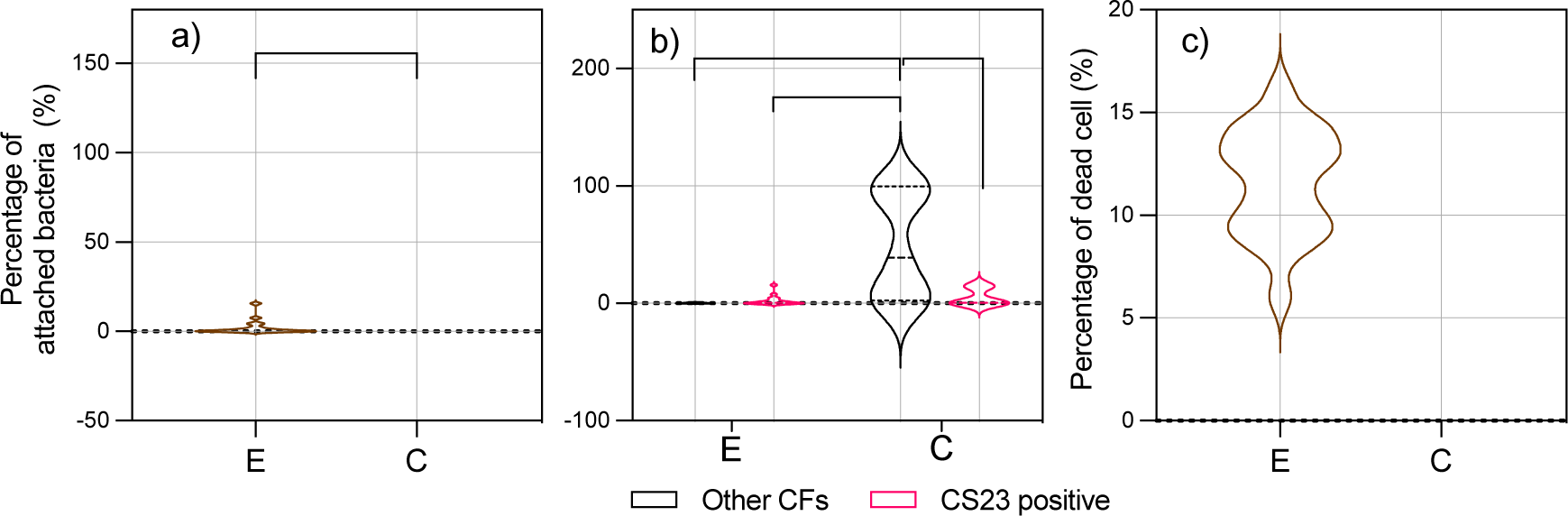
Bacterial adherence and cytotoxicity to human epithelial cells. **a)** Differences in the number of bacteria attached to Caco-2 monolayers according to the source of isolation (environmental and clinical) or **(b)** ETEC expressing CS23 vs. ETEC isolates expressing other CFs (non-CS23) of each group. C = clinical, E = environmental. Assays were performed in triplicate and the violin plots show the distribution of the data and the mean and standard deviation. The asterisks indicate a significant difference in the unpaired t-test (*P < 0.05 and ** P < 0.01) using GraphPad Prism 8.00. **c)** Cytotoxicity to Caco-2 cells infected by environmental and clinical ETEC isolates.

ETEC infection can disrupt/destroy the epithelial barrier, increase permeability, and induce an inflammatory response that damages intestinal cells (23). The evaluation of bacterial cytotoxicity showed that even though clinical isolates have a higher binding phenotype than environmental strains, both groups of strains were equally cytotoxic to the epithelial cell, as shown in Figure 3c.

### The circulation of ETEC isolates between the clinic and the environment is corroborated by PhenePlate typing

The affordability of DNA sequencing costs has improved, but the high cost of equipment, reagents, and limited resources in low- and middle-income countries (LMICs) present barriers to researchers accessing genomic technologies. Therefore, it is imperative to develop complementary, low-cost methods for detecting bacterial isolates. Here, we determine the bacterial relatedness of clinical and environmental isolates using a phenotypic screening method (PhenePlate) and compare its results to those obtained by whole genome sequencing. The PhenePlate method is a simple and rapid subtyping of *E. coli* based on comparing patterns in carbon source metabolism patterns and employs the UPGMA clustering method to describe the clonal relationship between bacterial isolates. As shown in Figure 4a, the 30 ETEC isolates were subjected to PhP typing and the resulting biochemical fingerprint of each isolate was compared and analyzed using the UPGMA clustering method (Table S3). This method identified seven types of PhP types (PhP type indicates that isolates with >0.975 similarities are considered identical and assigned a PhP type (P); see Methods) and seven single types of PhP (unique fingerprints) among isolates from our study (Figure 4a).

**Figure 4.**
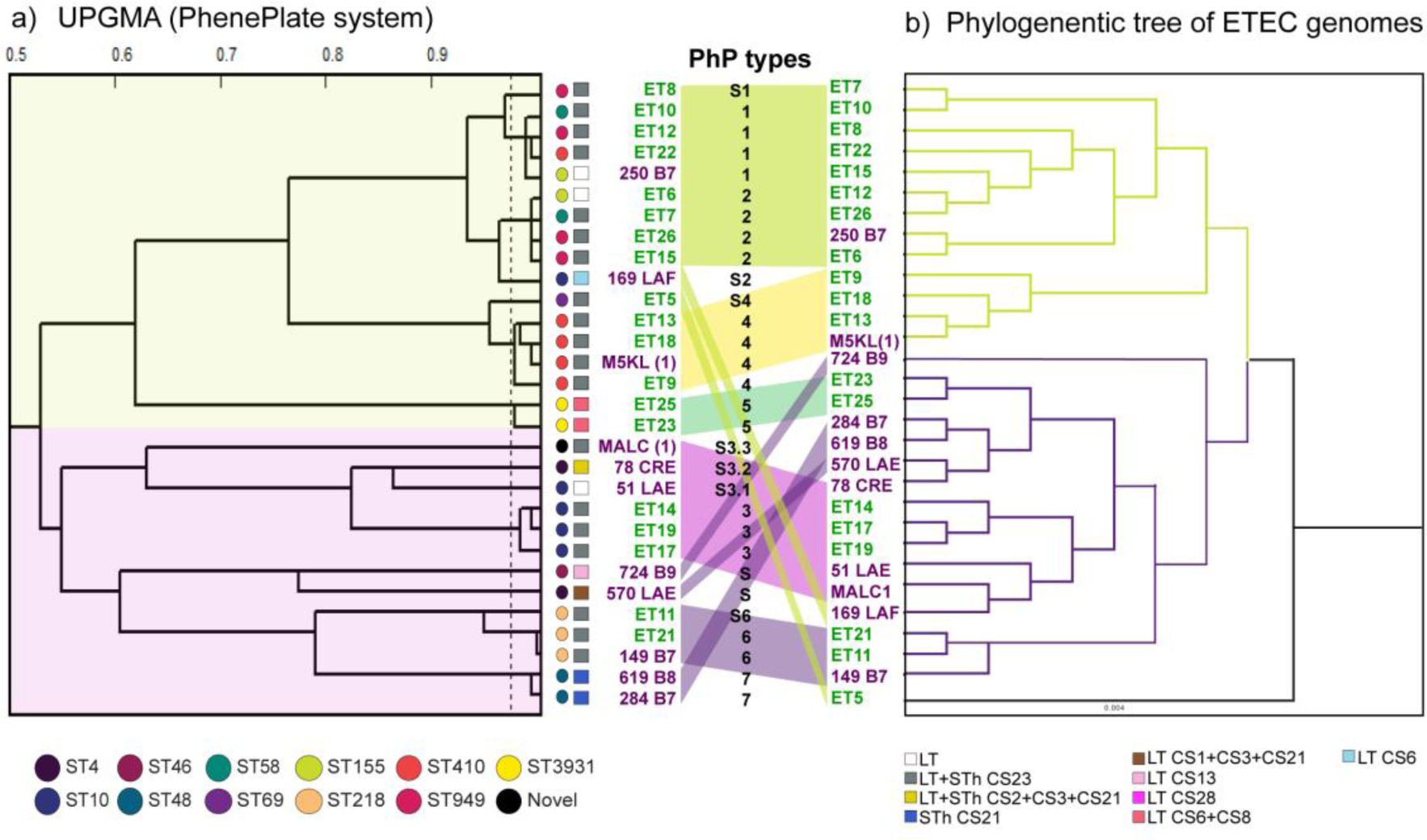
UPGMA dendrogram derived from clustering of the PhP types compared with the genome-based phylogenetic tree of the 30 ETEC genomes. **a)** UPGMA dendrogram derived from clustering PhP typing data from clinical and environmental ETEC isolates. The dotted lines indicate the levels of identity (correlation coefficient > 0.975). A colored link joins the same strains between two clustering methods. The major clusters were colored. **B)** Lineal tree modified from Figure 2a.

When comparing the UPGMA dendrogram and the phylogenetic analysis, we identified several congruencies in the topologies of both trees, and the PhP dendrogram was found to cluster the set of isolates in a similar manner as the WGS-based tree from Figure 2a. For example, cluster of the ST410 (M5KL, ET9, ET13 and ET18,) and ST218 (ET11, ET21, and 149 B7) genomes also that were clustered together and assigned distinctive PhP types (ST410:P4 and ST218:P6). These results corroborate the findings of genotypically and phenotypically highly similar isolates in both clinical and environmental settings. Another example is the P1 and P2 isolates which had a biochemical fingerprint highly similar (correlation coefficient > 0.9) and therefore clustered together. These two PhP types included the genetically close related STs *i.e.,* ST58, ST155, and ST949 from cluster A in Figure 2a. We also observed a correlation between other PhP types and ST groups, such as P3 (ST10), P5 (ST3931), and P7(ST48). Interestingly, isolates with single PhP type (S*n*) showed the most significant variation between the two methods. These data demonstrate that the PhP screening method, to a large extent, is consistent with the genomic resolution provided by the MLST technique. Hence, phenotypic fingerprinting is a powerful tool to detect differences in pathogenic circulating strains from various sources.

### Retrospective PhenePlate profiling of ETEC clinical isolates from endemic countries

Next, we tested the clonal relationship of our strains in context with the collection of ETEC isolates from endemic countries, including Bolivia, using the PhP method. The bacterial collection includes 152 ETEC strains of children (< 5 years) with acute diarrhea collected between 1980 and 2011. 106 of 152 strains were Bolivian strains obtained from several studies (10, 24) and the rest were from South (Argentina), Central (Guatemala) and North America (Mexico), Asia (Bangladesh and Indonesia), and Africa (Morocco and Egypt) (18, 25–27) (Table S4). In addition, reference ETEC strains of the main lineages were included in this collection (E925: L1, CS1+CS3+CS21; E1649: L2, CS2+CS3+CS21; E1441; L4, CS6+CS21; E1779: L5, CS5+CS6; E562: L6 CFA/I+CS21 and L7: E1373 CS6) (21). Of the 182 ETEC tested, 20 PhP types were identified, of which 7 (P1, P3, P8, P9, P11, P15 and P19) belonged to isolates obtained in this study (Figure 5, Table S5). These strains received new PhP types, and the former PhP assignment performed in Figure 4 is shown in parentheses. Most of the clinical and environmental isolates of this study were spread across the PhP dendrogram, forming their own clusters. For example, environmental CS23 ETEC strains P3 (also P2: ET6, ET7, ET26, and ET15) showed an identical PhP profile and therefore clustered with two other clinical Bolivian strains (CS21 E2381 and CF-neg E2404) and 1 Bangladeshi isolate (CS5+CS6 E5082). In this analysis, the PhP method placed strains that obtained unique PhP profiles from Figure 4 (724 B9 and 570 LAE) with P13 and P14 ETEC strains. The P13 group encompassed two more Bolivian CF-neg strains, while P14, in addition to the CS21 570 LAE strain, contained nine clinical strains of CS1+CS3+CS21. Another example is the environmental isolates expressing CS6+CS8 (E23 and E25) that had the same PhP profile as four other clinical Bolivian strains (CS14, CFA/I, and CF-neg) from 2009.

**Figure 5.**
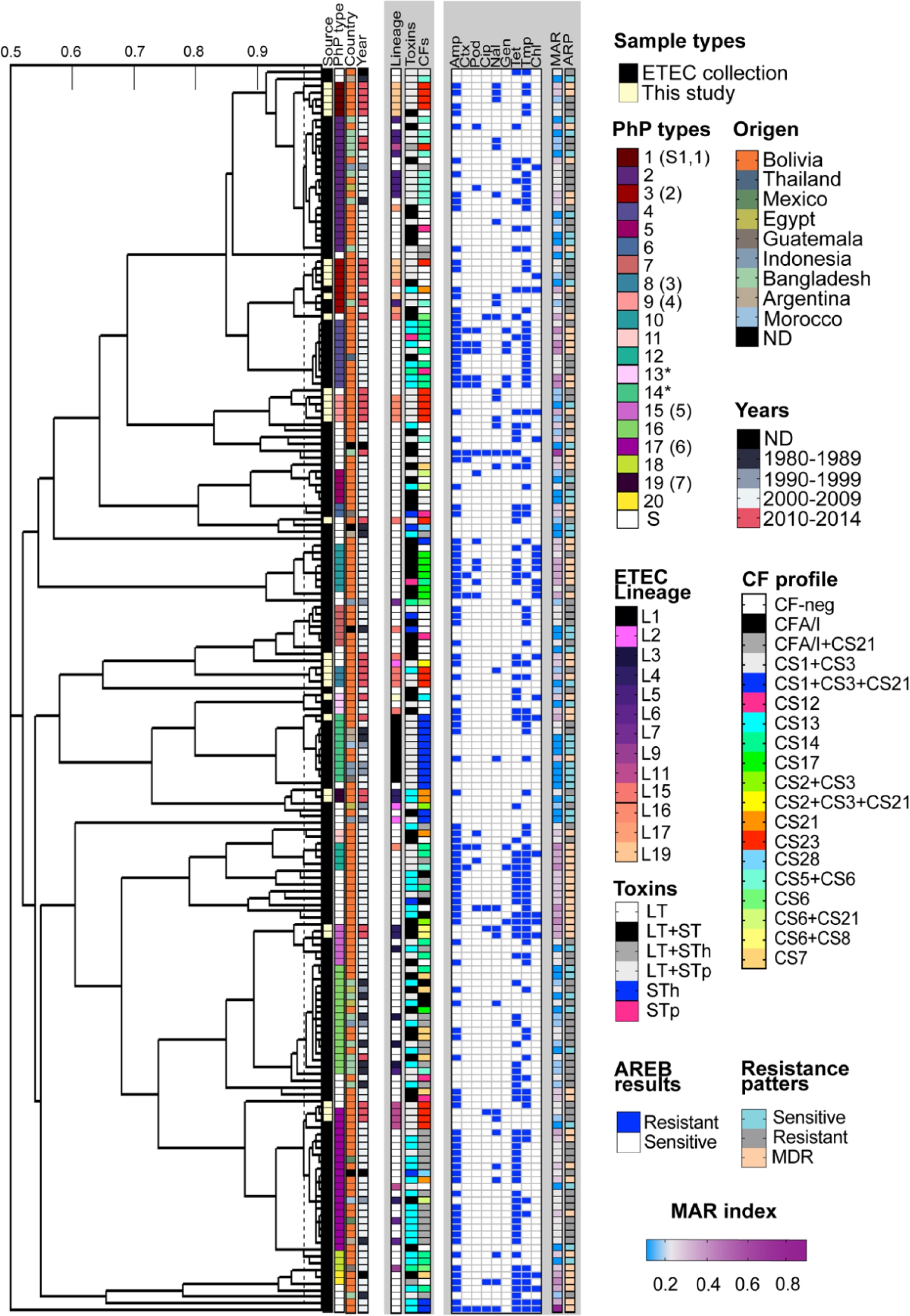
Comparison of phenotypic fingerprints from a collection of clinical Bolivian ETEC strains and isolates obtained in this study. The dendrogram shows the clustered PhP typing data, including the antibiotic resistance patterns obtained parallel by the AREB system. Information about the place and year of isolation, ETEC lineage, toxin, and CF profile is indicated in the legends in the figure. The identified lineage of ETEC isolates was included only for isolates of the ETEC collection included in the von Mentzer et al. (2014) study. Antibiotic resistance to the 9 AREB antibiotics, the multiple antibiotic resistance index (MAR), and antibiotic resistance patterns such as sensitivity, resistance, and multidrug resistance (MDR, ≤3 classes of antibiotics) were incorporated into the heatmap. The white coloration in the MAR index gradient indicates the critical limit of the MAR of 2. AMP, ampicillin; CTX, cefotaxime; POD, cefpodoxime; CIP, ciprofloxacin; NAL, nalidixic acid; GEN, gentamycin; TET, tetracycline; CHL, chloramphenicol.

There was also a notable correlation between the phylogenetic tree of 362 ETEC (Figure 1b) and the PhP cluster of Bolivian isolates (Figure 5). Our PhP data showed that the strains P1 and P3 characterized as members of the ETEC lineage L19 clustered with isolates P2 (CS5+CS6) and P4 (CS14 and CF-neg) isolates from the ETEC lineage L5 and L17, respectively. Slightly more distant in the dendrogram were found P9 (CS23) isolates that belonged to L16 and were linked to a group of mainly CF-neg isolates with unique PhP types and an uncharacterized lineage. According to von Mentzer’s study (18) and Figure 1b, the L19’s closest lineages were L20 (CS14, CS6, and CF-neg), L5 (CS5 + CS6), L17, (CS6, CS14, and CF-neg) and L16, which included isolates with diverse CF profiles, was more distant lineages. The L15 isolate 169 LAF was the only exception since the PhP method misplaced it with the L19 isolates. However, the L11 ETECs expressing CS23 that were circulating cross-settings (E21 and 149 B7) fell into the P17 cluster composed of distantly related L4, L6, and L9. Strains from these clusters were in their majority CFA/I+CS21 and isolated in Bolivia. In another example, in Figure 5, we found that strains P8 expressing CS23 (ET14, ET19, ET17) from lineage L15 clustered with other Bolivian clinical strains with a different type of PhP (P7) but the same lineage (CF-neg strains). The P14 and P19 strains, including L1 and L4 isolates, were the closest to the L15 strains. According to the tree in Figure 1b, L1 is closely related to L15 (Figure 5, Table S5).

Overall, the results of the PhP method in this extensive ETEC collection have shown that some, but not all, strains of this study shared similar metabolic fingerprints from clinical isolates collected previously in Bolivia and other endemic countries. Like genomic results, the population structure of CS23 ETEC based on the PhP method also shows high levels of complexity, extending along several PhP types throughout the dendrogram. Although only 45,4% (69) of the isolates had identified their ETEC lineage, the PhP dendrogram resembled well the population structure of the ETEC clades based on genomic data here (Figure 5, Table S5) and in the von Mentzer study (18). These data reinforce the notion of stable combinations of chromosomes and plasmids across the ETEC lineages, which seems to display conserved metabolic fingerprints. Further addition of more well-characterized representative ETEC strains will be needed to strengthen the discriminatory power of the PhP method.

### Antimicrobial resistance in ETEC strains during the past 12 years

Only ETEC strains previously isolated in Bolivia (104/152 strains) and strains obtained in this study were included for the screening of phenotypic resistance to antibiotics against nine antibiotics using the AREB method (see Methods). As described in Table 3, the total percentage of resistant, sensitive, and MDR strains as well as the MAR index between isolates from this study and the retrospective collection of Bolivian strains did not generate statistical differences. Only the comparison of the MAR index per year of isolation showed a statistically significant increase from 2007 to 2008 (ANOVA, P = 0.0004; Figure S4). Although resistance to ciprofloxacin and nalidixic acid has increased since 2009, the number of resistant strains remains low. These results showed that although resistance to clinically relevant antibiotics did not increase significantly over the past decade in this set of strains, almost 50% of ETEC isolates showed MDR.

**Table 3.**
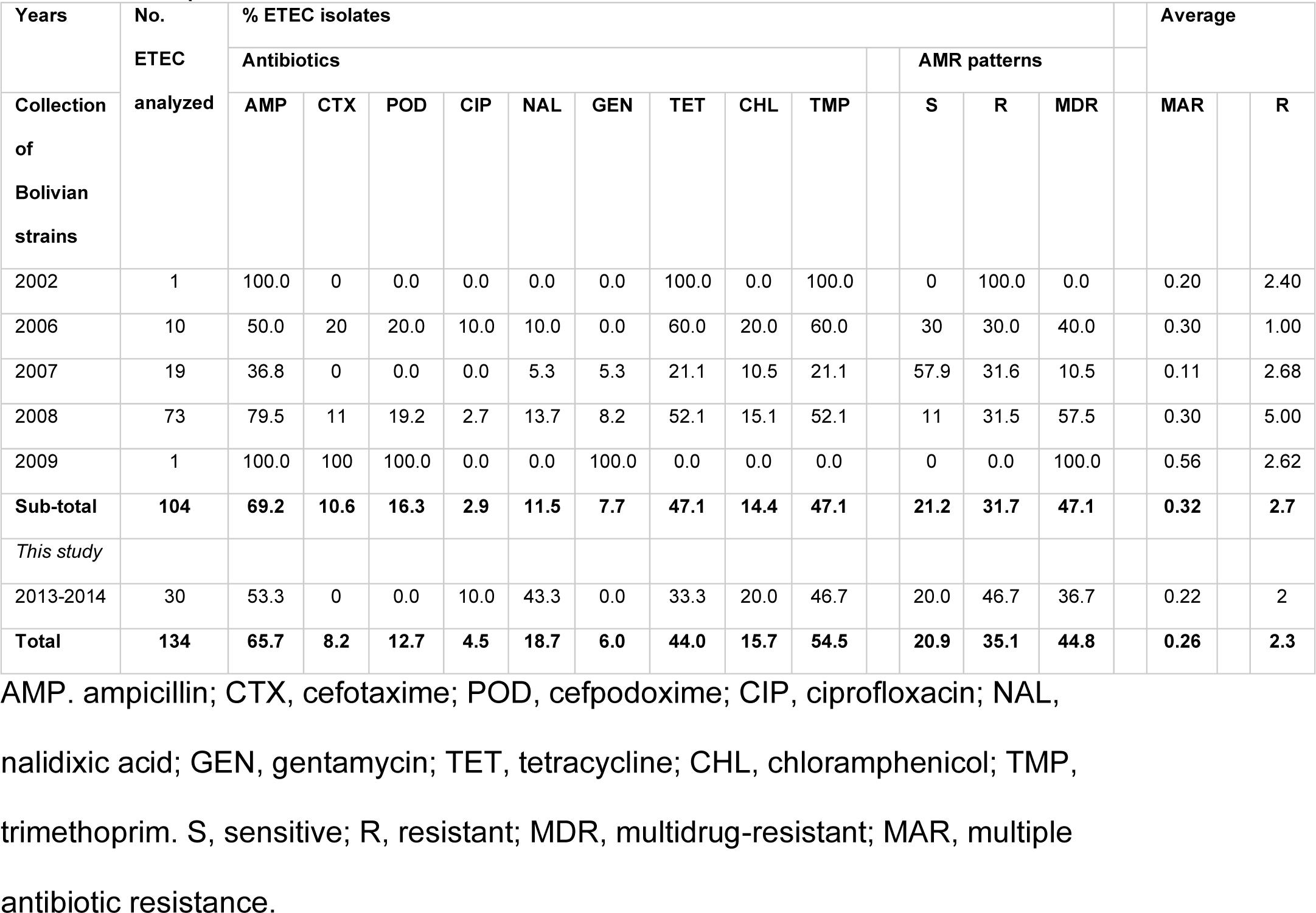
Comparison of annual resistance rates of Bolivian ETEC isolates.

## Discussion

Monitoring the dissemination of diarrheal pathogens between the human-environment interphase and the characterization of their antibiotic resistance patterns provides valuable information on the potential risk of outbreaks and treatment options. Monitoring is specifically relevant in countries where diarrheal diseases are endemic and socioeconomic factors such as inadequate clean water, poor sanitation (1), limited surveillance, and scarcity of microbiological laboratory infrastructure and genomic sequencing contribute to the burden of diseases and exacerbate the problem of AMR (28). Studies carried out in the region covered in this study have focused on characterization of virulence and antibiotic resistance in clinical isolates of ETEC (24, 29, 30), but mounting evidence suggests frequent detection of ETEC in drinking and environmental water (11, 14, 31, 32). Here, we use whole genome sequencing and the PhenePlate phenotypic screening assay to comprehensively track ETEC isolates from surface water and diarrhea. This study is the first of its kind in Bolivia, as our results provide evidence of transmission through the environment and clinical settings by specific clones of ETEC that were positive for the colonization factor CS23. The findings that most of the environmental isolates collected possessed the virulence profile LT + STh CS23 suggest that the Choqueyapu River could act as a reservoir for these types of bacteria and indicate that CS23 should be included in future and retrospective epidemiological studies. The high diversity in the virulence profiles among clinical ETEC compared to the ETEC of the Choqueyapu River could suggest that these strains may have other reservoirs or transmission routes.

It has been shown that *E. coli* can survive and persist in the environment for extended periods (33–36), and ETEC has been shown to remain viable for several months with preserved gene expression of virulence factors in water (37), highlighting that water is not only a reservoir but also a transmission vehicle (30). The waters of the Choqueyapu River flow from the pristine highlands through the city of La Paz, and further downstream, their waters reach the agricultural region, which is often used to irrigate crops and vegetables (12). Given that La Paz lacks a wastewater treatment plant and that the drainage of households, industries, and hospitals is discharged into the Choqueyapu River, it was not surprising to detect ETEC in the Choqueyapu waters, and our results correlate with other studies in this river that successfully isolated and identified ETEC and other diarrheagenic pathovars resistant to multiple antibiotics (11, 12, 31, 38, 39). Not only do the river(s), including the water sites selected for this study, have high levels of bacterial pollution (3.0×10^5^ ± 3.77×10^5^ MPN) that exceed the WHO guidelines for unrestricted irrigation water (WHO, 2016, *i.e.,* <1000 MPN/100 ml)(12) but also high levels of pharmaceutical pollutants (incl. antibiotics such as sulfamethoxazole) (15). Our study, in conjunction with other studies, highlights the high risk that the Choqueyapu River poses to the health of the community due to the transmission of pathogens by water and food. Therefore, the alarming levels of antibiotic residues present in its water could contribute to the selection and spread of resistant pathogens in sediments and waters and exacerbate the local and global AMR crisis.

ETEC is an important etiological agent associated with childhood diarrhea in Bolivia (10). Previous studies on the molecular epidemiology based on PCR detection of the most prevalent CFs (CFA/I, CS1-8, CS14, CS17, CS21) (24, 29, 40, 41) showed that CFA/I, CS14, and CS17 are the most common CF among diarrhea-associated ETEC strains; although 30-40% of all ETEC strains remained negative for any CF. In the last 10 years, research on CF-neg strains has led to the discovery of new CFs, including CS23 (42). Integrating whole genome sequencing provided a better resolution for the characterization of CFs and allowed the identification of non-traditional CFs from bacterial genomes, often neglected by surveillance and multicenter studies (17). The WGS of 125 ETEC isolates from diarrheal cases in Chile showed that, in addition to the prevalence of CFs similar to previous global studies, the novel CF CS23 was present in 4.8% of the isolates and indicates that this type of ETEC circulates in the region (43). With our WGS-based CF identification, we identified not only the CS23 operon in 19 ETEC isolates, but also five novel variants of this colonization factor and four genomes (51 LAE, 250 B7 (1), 724 B9 (1) and ET6) harboring a gene cluster encoding potential chaperone-usher (CU) assembled-pili. Interestingly, high-risk ST410 in both clinical and environmental samples consistently contained the same CS23 allele (AalE3). More functional to verify expression, assembly, and adherence patterns of the putative-CS23 and the putative CU-pili must be performed.

Interestingly, the genetic relationship of CS23 with other CFs has shown a close relationship with porcine F4 fimbria and other F4-like fimbria mediating adhesion to plant surfaces such as lettuce and crop leaves (4). All environmental CS23-positive ETEC isolates came from water samples collected near or on farmlands in the south of the city of La Paz, where legume, vegetable, and corn crops are grown and sold to retailers that sell the products in more than 80 markets in the cities of La Paz and El Alto. In 2016, a survey on the microbial safety of foods and vegetables in this area of La Paz confirmed the presence of high levels of bacterial pathogens and parasites and concluded that the health of farmers and consumers is directly and indirectly at risk (44). In this study, the Choqueyapu River has been shown to play a role as a reservoir and a route of spreading and transmission to the local population.

The population genomics of clinically relevant ETEC isolates from indigenous travelers, and asymptomatic patients have been extensively studied, demonstrating remarkable genomic diversity (18, 45–47) with distinct lineages with conserved plasmid-encoded virulence profiles (18, 21). Despite most diarrheal cases being linked to environment or food sources (30), environmental ETEC remain understudied. Our study provides a snapshot of the diversity and complexity of Bolivian ETEC genomes through a phenotypic and genomic approach and by integrating two settings of the One Health concept. Our phenotypic and genomic approach reveal a remarkable diversity of CS23 ETEC isolates in terms of biochemical fingerprints (PhP types), STs, and *E. coli* phylogroups (A, B1, C, and D). A comparison with the 362 globally distributed clinical ETEC genomes from von Mentzer’s shows that non-CS23 Bolivian isolates belong to established, clonally distributed ETEC lineages (L1-L5) with a conserved combination of toxins, virulence factors and chromosomal background (18, 21). On the other hand, CS23 positive isolates from Bolivia span multiple linages, where mostly clinical isolates are CF negative. Hence, this suggests that the presence of CS23 in both environmental and clinical isolates may not indicate a clonal outbreak but rather reflects the diverse origin and range of habitats of the bacteria (48).

Analysis of the longest contig (89,895 bp) containing the CS23 gene cluster shows that the plasmid background is conserved, like other CFs (21). Interestingly, a pattern is evident between the chromosomal background and the CS23 AalE variant (Figure S1). Whether or not the plasmids harboring the CS23 gene cluster are conserved requires additional analysis. Furthermore, the location of CS23-positive ETEC in the terminal branches of the phylogenetic tree of representative ETEC genomes could suggest a recent acquisition (after 2011) of mobile genetic elements, *i.e.,* plasmids that contributed to a faster diversification of these strains between different lineages, as observed in *E. coli* genomes from animal and environmental sources (48).

Our study has provided evidence of the spread of ST410 and ST218 LT+STh CS23-ETEC strains between hospitals and the environment. The identification of identical clones with the same CS23 alleles in both stool samples and the Choqueyapu River supports the hypothesis of fecal contamination of the river as the cause of spread. This could have occurred through hospital or hoursehold discharges into the river. The association of CS23 with porcine or palnt-derived virulence factors also raise the possibility of contaminated vegetables or meat as potential vehicles for the transmission to the local population.

In previous studies, it has been shown that ETEC outbreaks can occur due to contaminated salad vegetables (49–51). Colonization factors such as CS21 or virulence factors, such as flagella (52) and the *etpA* filaments (52) play a role in adhesion to food-related surfaces. Our findings that the majority of CS23-expressing ETEC strains were strong biofilm producers suggest that biofilm formation may also play a role in colonization and persistence of these bacteria (53) in sediments, crops, or fecal matter. The Choqueyapu River is also known to be contaminated with biofilm-inducing pollutants such as sub-MIC concentrations of antibiotics (54, 55) and offer the ideal conditions for growth and biofilm production with its constant flow providing the oxygen and nutrients (56). On the basis of these observations, we hypothesize that during dry seasons, bacterial accumulate in biofilms only to detach and spread to other locations or hosts during the rainy season when water flow rates are high. These bacteria then re-enter the human host through contaminated water or food.

Ingested ETEC strains must adhere to intestinal cells to secrete toxins and initiate infection (1). Here, we demonstrate that clinical ETEC isolates have a superior adherence to CaCo-2 cells, compared to environmental ones. The greater attachment ability of clinical isolates, particularly those expressing other CFs besides CS23, can be attributed to their previous colonization and infection of the human host. However, contrary to our results, F. Del Canto et al. (4) demonstrated that CS23 ETEC had a higher adhesion capacity than the CFA/I ETEC strain H10407. The same study revealed that the low levels of adherence seen in certain colonies of a CS23 ETEC strain were due to spontaneous mutations in the major fimbrial structural subunit AalE, which plays an essential role in the adherence of ETEC. The natural polymorphism identified in the AalE gene of our CS23 ETEC isolates could have negatively impacted the bacterial adhesion; however, further research is required to understand the role of the AalE gene in bacterial adhesion. Despite some environemtal isolates possibly losing some adhesion properties *in vitro* infection models, they still possess virulence properties according to our cytotoxicity assays.

ST410 is a high-risk MDR clone of *E. coli* commonly associated with carbapenemase genes that carry *bla*_OXA-181_ and *bla*_NDM-5_ and with cross-sectorial transmission between wildlife, humans, companion animals, and the environment (57, 58). ST410 was previously reported in soil samples from the Choqueyapu River and carried the *bla*_OXA-1_ gene (11). Studies have also found that *E. coli* ST410 is associated with vegetables on the municipal market in Ecuador (59) or surface water from Mexico’s agricultural drainage (60). The ST410 isolates described in this study were not MDR, but carried the CS23 operon in combination with LT and STh. A study that has assessed the transfer of multidrug resistance plasmid from Choqueyapu’s waterborne bacteria to *E. coli* indicated that the IncN plasmid carrying a wide range of ARGs and *intl1* were transferred with high frequency (61). A potentially dangerous scenario is in which virulence genes and last-generation MDR genes are combined in one ST410 isolate. Since CS23-positive isolates were detected in children’s diarrheal stool and environmental water, such combinations could promote an emerging diarrheagenic clone that can spread more easily. The apparent ability of ST410 to form biofilm at GI and environmental temperatures could further facilitate survival and spread in different niches (62).

The high levels of MDR among ETEC isolates reported in this study agree with other reports in Bolivia (10, 29, 40), Latin America (63), other LMICs (64–66) as well as in other diarrheagenic *E. coli* (19) and reflect the high rates of empiric prescription, often not corroborated by clinical microbiology laboratories, self-medication, and lack of regulation of antibiotic use observed among LMICs. Although low levels of quinolones (ciprofloxacin and nalidixic acid) were reported in this study, our retrospective evaluation and comparison with previous studies indicate an increasing trend over time of ETEC resistant to fluoroquinolones in Bolivia. This increasing trend was also reported among isolates of ETEC and EAEC causing traveler’s diarrhea (67) and *Shigella* species associated with diarrhea (68). The burden of antibiotic resistance on ETEC associated diarrhea in children can be significant, particularly in settings where antibiotic resistance is prevalent, leading to more severe or persistent diarrhea and complicating the use of prophylactic antibiotics (69). Hence, it is an important global health issue that requires more attention, funding, capacity building, research, and development (70).

The use of typing methods for the accurate discrimination of different bacterial isolates of the same specie but different origen is crucial for effective infection prevention and control. We evaluated the performance of a fast and cost-effective phenotypic detection method based on the ability to ferment various sugars, compared to the genome-based MLST typing method. Our results indicate that the PhP assay is a reliable and useful tool for studying the bacterial population of coliforms in various sources, such as water (71–73) and intestinal flora (74, 75), as in previous studies.

Additionally, our findings support the nation that the PhP assay provides results comparable to those of MLST-type resolution and reveals conserved metabolic fingerprints associated with specific *E. coli* sequence types (STs). This was demonstrated in our earlier studies of water pump station samples in Norway (71–73), and now we have extended this concept to the ETEC lineages, which showed conserved metabolic fingerprints. The use of the PhP assay, which can identify MDR pandemic clones ST131, ST648 and the high-risk clone ST410, makes it a useful screening tool in resource-poor settings where sequencing multiple is unaffordable.

The association between ETEC expressing CS23 and diarrhea can only be speculated on due to the low number of samples analyzed. Furthermore, a specific metabolic fingerprint could not be identified as the level of diversity in the ETEC isolates expressing CS23 was high and the resolution of PhenePlate was low.

In conclusion, we report that ETEC isolates positive for CF CS23 are common in environmental water in Bolivia, and strains carrying this CF circulate between the environment and human hosts. CS23 could be more common than previously anticipated and its inclussion in the monitoring of ETEC epidemiology and analysis of its virulence protential is crutial. Our study provides a framework for tracking bacterial pathogens using the PhenePlate that can be used across all components of One health (humans, animals, and the ecosystem) due to its high-throughput system, feasibility, and low cost. The integration of this low-cost tool with genomic technologies will allow for efficient monitoring and detection of emerging pathogens in areas where surveillance is minimal and data sparse.

## Materials and Methods

### Isolation of strains

#### Environmental isolates

The study area comprises different locations along the Choqueyapu River. This river crosses the city of La Paz, Bolivia, and is highly polluted due to the discharge of domestic and industrial wastewater (12). Surface water samples were collected from five locations downstream of the Choqueyapu river. The sampling sites included five communities: Palomar, Mecapaca, Lipari, Aranjuez, and Carreras. They are characterized by agricultural areas that use Choqueyapu waters to irrigate vegetable crops. Water samples (300 ml) collected in sterilized bottles and stored at four °C were processed on the same day of the collection. The water samples were filtrated through 0,45 µm pore-sized cellulose membrane filters (Millipore Sigma Aldrich, St. Louis, MO) with a vacuum/pressure pump system (Pall Life Sciences, Ann Arbor, MI). For enrichment, the filters were incubated overnight in EC Broth (Oxoid, Basingstoke, Hampshire, England) at 37°C. The enriched EC broth cultures were seeded on MacConkey agar (Oxoid, Basingstoke, Hampshire, England) and incubated overnight at 37°C. From each water sample, a total of twenty *E. coli*-like lactose-positive colonies were selected and subjected to PCR to detect ETEC toxins (LT, STh, and STp) as previously described (29). The reference ETEC strain H10407 was used as a positive PCR control. LT and/or ST-positive isolates were considered ETEC and cryopreserved (LB-glycerol, 85:15%, v/v) at −80°C for further experiments.

#### Ethical statement

Since the present study was approved as part of routine diagnostic practice within the Diarrheal Disease Project and the National Rotavirus Surveillance program in Bolivia (10), it was not necessary to obtain specific approval from the respective hospital ethics committees or informed consent from patients. No names or personal ID were included, and it was not relevant for the study.

#### Clinical isolates

Identification of ETEC isolates from fecal samples collected from children under five years of age with acute diarrhea attending Los Andres Hospital (El Alto city) and Materno Infantil Hospital (La Paz city) was carried out as described by Gonzáles et al. (24). The clinical isolates were stored in LB with 12% glycerol at −80C at the Institute of Molecular Biology and Biotechnology (IBMB) until further evaluations. The frozen strains were grown overnight in EC broth and then placed on MacConkey agar plates. The isolates were re-evaluated for ETEC virulence factors by PCR using primers targeting ETEC toxins and CFs (24).

### DNA extraction, whole genome sequencing, assembly, and annotation

Genomic DNA was extracted as previously described with standard methods (18) using the DNeasy blood and tissue extraction kit (Qiagen) and eluted in 200 ul of MilliQ water. DNA concentration was measured using Qubit, and 50 ng of DNA was used for library preparation. Sequencing libraries were prepared using the MGI FS library prep set according to the manufacturer’s instructions. The TapeStation D1000 kit (Agilent) was used to evaluate library quality. Circularized DNA of equimolarly pooled libraries was prepared using the MGI Easy Circularization kit (MGI Tech). The DNBseq 2×100 bp paired-end sequencing was performed using a DNBSEQ G400 instrument (MGI) according to the manufacturer’s instructions. Raw sequencing reads were assembled using the Velvet assembler.

Annotated assemblies were produced using the pipeline described in (76). For each sample, sequence reads were used to create multiple assemblies using VelvetOptimiser v2.2 and Velvet v1.2.10 5 (77). An assembly improvement step was applied to the assembly with the best N50, the contigs were scaffolded using SSPACE v2.0 (78) and the sequence gaps were filled using GapFiller v1.11(79). Automated annotation was performed using PROKKA v1.5 (80) and a genus-specific database (’Escherichia’) from RefSeq (81). All the software developed by Pathogen Informatics at the WSI is freely available for download from GitHub [Pathogen Informatics, WSI, https://github.com/sanger-pathogens/vr-codebase, Bio-Assembly-Improvement: Improvement of genome assemblies by scaffolding and gap-filling, Pathogen Informatics, WSI, https://github.com/sanger-pathogens/assembly_improvement] under an open source license, GNU GPL 3. The pipeline improvement step is also available as a standalone Perl module from CPAN (http://search.cpan.org/~ajpage/).

### Genomic profiling: virulence, phylogroups, and multilocus sequence type

The genomic toxin and CF profiles were determined in two steps, 1) running Abricate using a custom database (https://github.com/avonm/etec_vir_abricate) and 2) BLASTn of genomes against the custom database (https://github.com/avonm/ETEC_vir_db) using the in-house script that creates a blast file that can be loaded into the Artemis Comparison Tool (ACT) (https://www.sanger.ac.uk/tool/artemis-comparison-tool-act/) along with the reference database and the annotated genome of interest. The phylogroup of ETEC strains was determined using ClermonTyping (v20.03) (82). The multilocus sequence type (MLST) of Bolivian isolates was predicted using ARIBA (v2.14.6) (83) following the Achtman scheme and the database ‘Escherichia coli #1’ with the reads as input data.

### Phylogenetic analysis

The core genome was defined using Roary (84) with the split-paralogs option switched off and using mafft as the aligner (command: roary -p 20 -g 60000 -e -n -s -f output_dir *.gff). SNP-sites (85) was used to extract SNP sites. Furthermore, the number of constant sites for each nucleotide (ATGC) was also determined using SNP-sites. The relationship between Bolivian ETEC isolates and in context with the 362 previously published ETEC isolates (18) was investigated (Fig 1a and 1b). Estimated maximum-likelihood phylogenies were generated with IQTree (v1.6.10) using the GTR+F+I model and 1000 ultra-fast bootstraps (-bb), and the constant sites from SNP-sites were included for linear scaling of the branch lengths (-fconst) (command: iqtree -s aln_file-mem 4G -nt 4 -bb 1000 m GTR+F+I -fconst).

The phylogenetic trees, along with metadata, were visualized using R (v4.0.2, 2020-06-22, URL: https://www.project.org/), explicitly using the R packages GGTREE (86) and GGPLOT2 (v3.3.2, URL: https://ggplot2.tidyverse.org).

### Genomic analysis of CS23 gene clusters identified in Bolivian genomes

CS23 positive isolates were analyzed using BLASTn against a reference database of known CFs, including CS23, and visualized in ACT. The gene clusters encoding CS23 were then manually extracted and individual genes were used as input for an all-vs-all BLASTn together with the F4 and CS23 references (https://github.com/avonm/ETEC_vir_db/blob/main/ etec_vir_master.fasta). The structural major subunit AalE of all CS23 positive isolates was extracted and aligned using mafft (87). The aligned sequences were used as input for IQTree (model: WAG+F+G4) (http://www.iqtree.org/) to build a phylogenetic tree. In total, five variants of AalE were evident, which was corroborated by pairwise alignment in Jalview of each AalE variant.

### PhP and AREB analysis

A rapid, semiautomated, and computerized typing method was used for *E. coli* based on measurements of the kinetics of 12 biochemical reactions, known as the PhenePlate system (PhP-RE) for typing. ETEC isolates were obtained from this study, and Bolivian and worldwide ETEC collections were subjected to the PhP assay (PhP PhenePlate (PhP) typing PhPlate Microplate Techniques AB, Saltsjö-Boo, Sweden; http://www.phplate.se) as described elsewhere (73, 88). Briefly, pure *E. coli* colonies were extracted from the agar using sterile tooth sticks and inoculated into PhP-RE plates, previously filled with 150 µl of PhP suspension medium. Ten µl of the bacterial suspensions were transferred to the wells of all other columns of the PhP-RE plate. Simultaneously, 20 µL of the initial bacterial suspension of the PhP-RE plate was transferred to the first column of AREB plates containing ten antibiotics [column 2-11: ampicillin (32µg), cefotaxime (2µg), ceftazidime (16 µg), chloramphenicol (32µg), ciprofloxacin (4µg), gentamicin (16µg), nalidixic acid (32 µg), cefpodoxime (3µg), tetracycline (16µg) and trimethoprim (16µg)] and filled with 200 µl of iso-sensitest broth (Oxoid), then 10 µl of suspensions were dispersed in each column (2–12) of AREB plates. The plates were incubated at 37°C for 18-24 h, and images of each plate were produced using a desktop scanner (HP G4050). Using PhenePlate™ software (PhPlate Microplate Techniques AB), images of the PhP-RE plate were transformed into absorbance data to generate the biochemical fingerprints of all isolates and, compared to each other, the similarity between each pair and create dendrograms. The reproducibility of the assay was used to determine the identity level (ID). The ID level was defined as the mean correlation coefficient between multiple assays of the same isolates minus two standard deviations (SD), corresponding to a confidence level of 95%. Isolates with a correlation coefficient higher than the ID level were considered identical and assigned to the same PhP type. The single PhP types (Si) contained only one isolate. The software also compared the amount of bacterial growth in each well of the AREB plate against the control well for the same bacterial isolate (column 12) and printed out resistance rates as 0 (no growth, susceptible), 1 (weak response, the result was controlled by visual inspection) and 2 (similar amount of growth as in the control well, resistant). For the final analysis, weak growth was considered to indicate resistance. The total antibiotic resistance in a population was measured using the multiple antibiotic resistance index (MAR), calculated as the mean proportion of resistance for isolates. The maximum possible MAR value is 1.00, obtained when all isolates are resistant to all antibiotics tested (89, 90).

#### Antimicrobial susceptibility test

The sensibility to antibiotics not included in AREB plates was determined using the disk diffusion method according to EUCAST guidelines (https://www.eucast.org/clinical_breakpoints/). The following antibiotic disks were tested: azithromycin (15µg), amikacin (30µg), tobramycin (10µg), streptomycin (10µg), tigecycline (15µg), and trimethoprim-sulfamethoxazole 1:19 (25µg). *Escherichia coli* ATCC® 25922 and *Staphylococcus aureus* ATCC® 25923 were used as reference strains. Strains that exhibit resistance to one or more agents in the last three different antimicrobial categories are considered multidrug-resistant (MDR).

#### Biofilm formation

As previously described, bacterial strains were tested to assess *rda*r morphotype formation (91). The bacterial strains were grown on LB-no salt agar (LBns) overnight at 37°C and resuspended in PBS 0.001 pH 7.4 (Sigma). The bacterial suspension was adjusted to an OD_600nm_ of 3, and 5 μl was spot inoculated onto LBns supplemented with Congo red (40 μg/ml) and Coomassie brilliant blue (20μg/ml). The colony morphology was observed after incubation at 37 ° C or 28 ° C for 48 hours; photographs were taken after incubation and the diameter was measured.

### Cell culture assays and infection

Caco-2 cells (human colon adenocarcinoma cells) were obtained from the American Type Culture Collection and were grown in Dulbecco’s modified Eagle medium (DMEM, Sigma-Aldrich) supplemented with 10% (v/v) fetal bovine serum, 1% streptomycin-penicillin, and 1% non-essential amino acids. Cells were grown in T-25 cell culture flasks at 37°C with 5% CO_2_ in an atmosphere of 5% CO2/95% air with constant humidity. Confluent cell culture flasks were trypsinized and subcultured when necessary. For bacterial infection experiments, cells were seeded at a density of 4000 cells/cm2 in 24-well tissue culture plates containing treated glass coverslips to grow adherent cells. Clinical and environmental ETEC bacterial isolates previously characterized, were grown overnight in LB agar plates at 37°C one day before infection. The same day of infection, bacterial strains were resuspended in sterile PBS, adjusting to a desired OD. Cell culture wells containing growing Caco-2 cells were washed three times with sterile PBS. For infection experiments, DMEM medium without antibiotics was used with 10% heat-inactivated fetal bovine serum (30 min at 56°C). Caco-2 cells were infected with previously prepared bacterial suspensions at a MOI of 200: 1 for an incubation time of 3 h at 37 ° C and a 5%CO2 atmosphere. Each bacterial strain was tested in triplicate.

#### Adherence assay

Infected Caco-2 cells were washed three times with sterile PBS to remove non-adherent bacteria. The Caco-2 cells were then fixed with methanol and dyed with Giemsa stain. The stained glass coverslips were removed from the culture wells and mounted on a microscope slide (three glass coverslips per strain). The adhesion of each bacterial strain was measured by counting the number of adherent bacteria in a total of 20 microscope fields (magnification 1000X). The number of infected and non-infected Caco-2 cells in the same microscope fields was also registered for further analysis.

#### Cytotoxicity assay

The live/dead viability/cytotoxicity kit for mammalian cells (Molecular Probes, Eugene, OR) was used to assess the cytotoxicity of ETEC bacterial strains in Caco-2 cells. Briefly, after Caco-2 cell infection with bacterial strains (described above), cell culture wells were washed three times with sterile PBS and cells were incubated with 2 µM calcein AM and 4 µM ethidium homodimer-1 for 30 min at room temperature. After incubation, stained glass coverslips were removed from the wells and mounted on a microscope slide in triplicate per strain. Microscope slides were observed under a fluorescence microscope (Leica DM LB2, Wetzlar, Germany), where viable cells showed a green fluorescent color and non-viable cells showed a red fluorescent nucleus. Viable and non-viable Caco-2 cells were counted and registered for further analysis.

### Data availability

The complete sequences of ETEC were deposited in the NCBI database within the BioProject SRP416785.

## Acknowledgments

We thank the clinical and laboratory team at the Instituto de Biología Molecular y Biotecnología, La Paz, Bolivia, and the sequencing team at the Centre for Translational Microbiome Research (CTMR) at Karolinska Institutet, Stockholm, Sweden. The authors also thank Ass. Prof. Åsa Sjöling for comments on the manuscript. This work was funded by the Swedish Research Council (EJ; grant 2019-04202 and AvM; grant 2018-06828), Swedish Research Links (EJ; 2017-05423) and Swedish Society for Medical Research (AvM).

## Supplementary Material

### Supplementary Tables

**Table S1.**Population characteristics, virulence factors, and genomic characterization of ETEC isolates included in this study. AD, acute diarrhea.

**Table S2.**Frequency of resistant ETEC isolates to 16 antibiotics tested by the disk-diffusion method. The P values were calculated using Chi-square and Fisher’s Exact test using GraphPad Prism 8.00. (**) Indicates statistical significance < 0,001. AMP, ampicillin; AZM, azithromycin; CIP, ciprofloxacin; NAL, nalidixic acid; GEN; gentamicin; AK, amikacin; TOB, tobramycin; STM, streptomycin; TET, tetracycline; TGC, tigecycline; CHL, chloramphenicol, TMP–SMX, trimethoprim–sulfamethoxazole; CTX, cefotaxime; POD, cefpodoxime; MER, meropenem and IMP, imipenem. Na; not applicable.

**Table S3.**Dendrogram derived from the UPGMA clustering of ETEC strains screened by the PhP method. Vertical dotted lines denote the pre-defined identity level. The total diversity (population diversity) was calculated using Simpsońs index of diversity among the isolates was 0.929 meaning high degree of diversity. Si, singleton (unique PhP profile). All data handling, including calculations of similarities, cluster analysis, and calculation of diversities, was performed using the PhPWIN software as described by Kuhn et al. (1). Values 0 – 30 represent the absorbance multiplied by 10 obtained after reading the test results using the desktop scanner. The PhP suspending medium contains the pH indicator bromothymol blue that changes to yellow at acid pH (bacterial fermentation) and blue at natural and alkaline pH.

(1) Kuhn, I., Brauner, A. and M Möllby, R. (1990) Evaluation of numerical typing systems for Escherichia coli using the API50 CH and the PhP-EC systems as models. Epidemiol Infect105, 521–531

**Table S4. List of 152 human ETEC isolates that were subjected to PhP typing.** Of the total of bacterial isolates, 41 isolates were included in von Mentzer’s study (1) and 38 of them were determined their ETEC lineage (L1-17). Geographic and temporal data and the virulence profile (colonization factors and toxins) were also included. ND, not determined.

(1) von Mentzer A, Connor TR, Wieler LH, Semmler T, Iguchi A, Thomson NR, Rasko DA, Joffre E, Corander J, Pickard D, Wiklund G, Svennerholm AM, Sjoling A, Dougan G. 2014. Identification of enterotoxigenic Escherichia coli (ETEC) clades with long-term global distribution. Nat Genet 46:1321-6.

**Table S5. Characteristics of the 182 ETEC isolates tested using the PhP method.** All isolates were coded, and the PhP fingerprint, AREB profile, year and country of isolation, virulence profile and ETEC lineage (colonization factors and toxins) are shown beside them. The PhP types calculated in Figure 4 are shown in parentheses. Sample type: 1, retrospective collection of ETEC strain collection; 2, isolates from this study; Country: 1: Bolivia; 2, Argentina; 3, Bangladesh; 4, Indonesia; 5, Guatemala; 6, Egypt; 7, Mexico; 8, Thailand. AREB: AMP, ampicillin; CTX, cefotaxime; POD, cefpodoxime; CIP, ciprofloxacin; NAL, nalidixic acid; GEN, gentamycin; TET, tetracycline; CHL, chloramphenicol.

### Supplementary Figures

**Figure S1:**
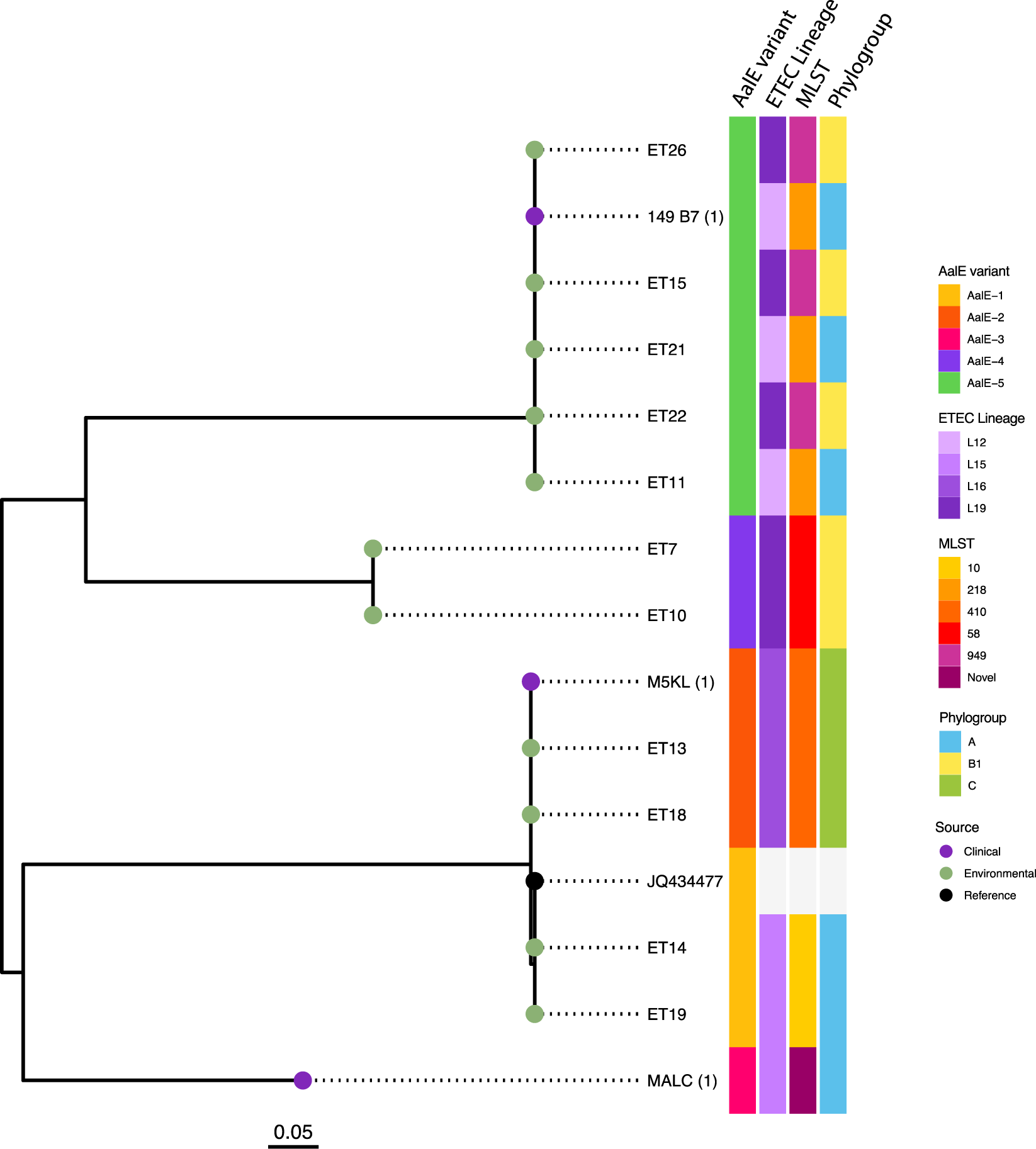
Maximum likelihood tree based on the amino acid sequence encoding the major subunit AalE in CS23 positive isolates. BLASTp comparison revealed five different variants of AalE (AalE-1 through 5). The ETEC lineage (based on the phylogenetic tree in Figure 1a and the predicted MLST and the phylogroup of each isolate are included in the heatmaps. The source of the isolate is color coded at the tips and the scale bar represents the substitutions per variable site.

**Figure S2.**
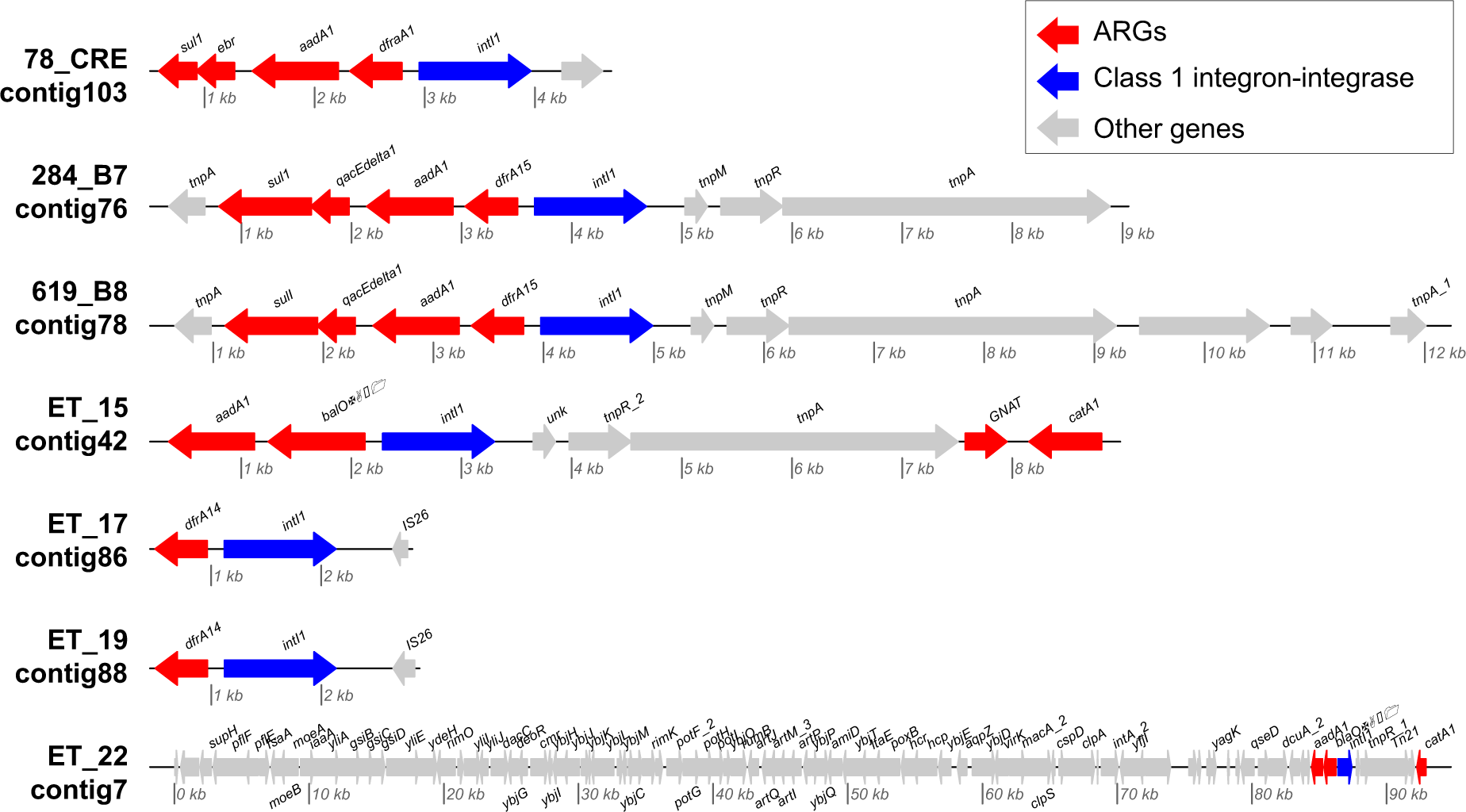
Gene organization of contigs carrying the class 1 integron-integrase gene *intI1* and ARGs. An arrow denotes the gene area. The ARGs, the intI1 gene, and other features are colored.

**Figure S3.**
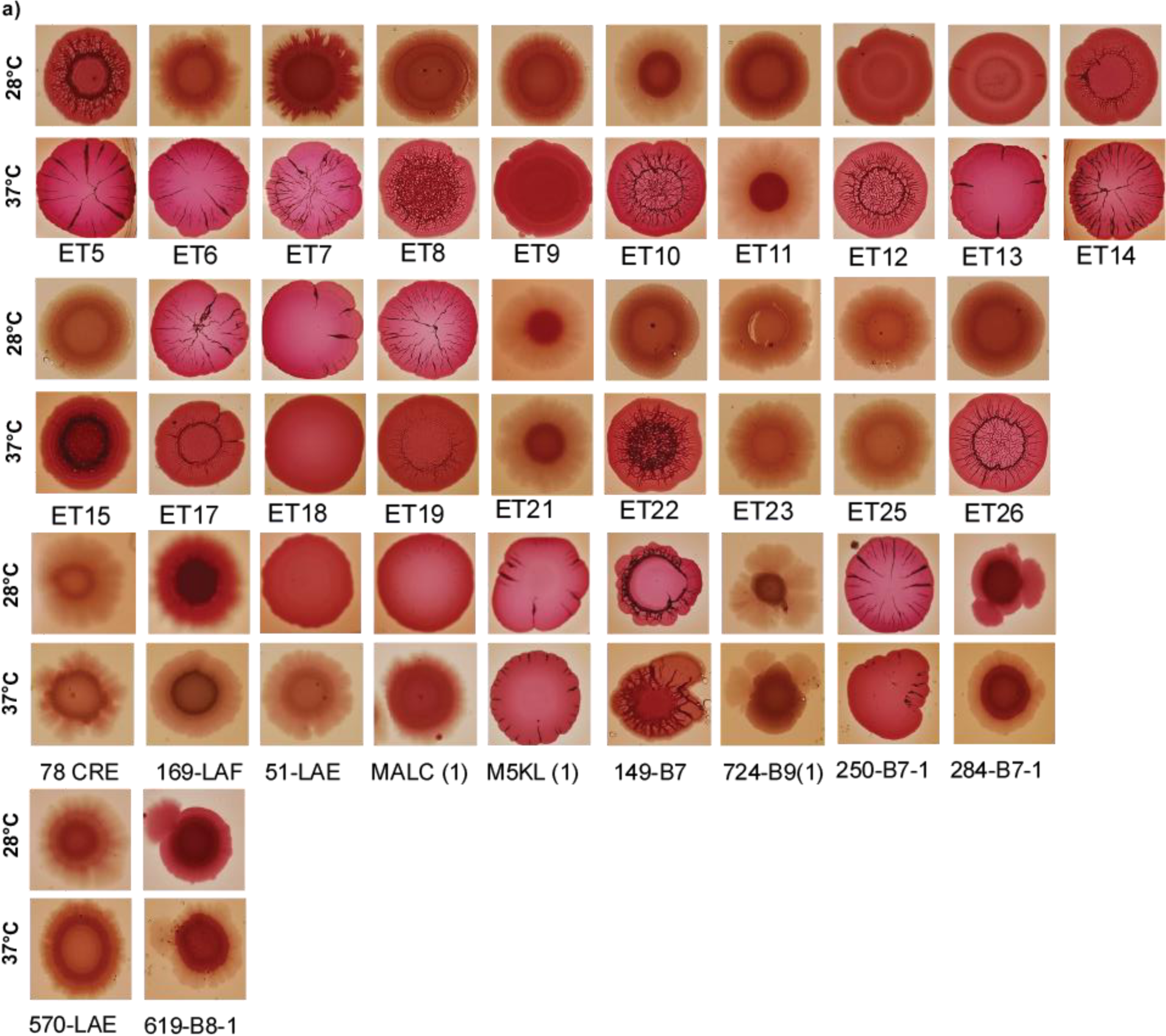
Effect of temperature on the formation of morphotypes in clinical and environmental ETEC isolates at 20 ° C and 37 ° C for 48 h.

**Figure S4.**
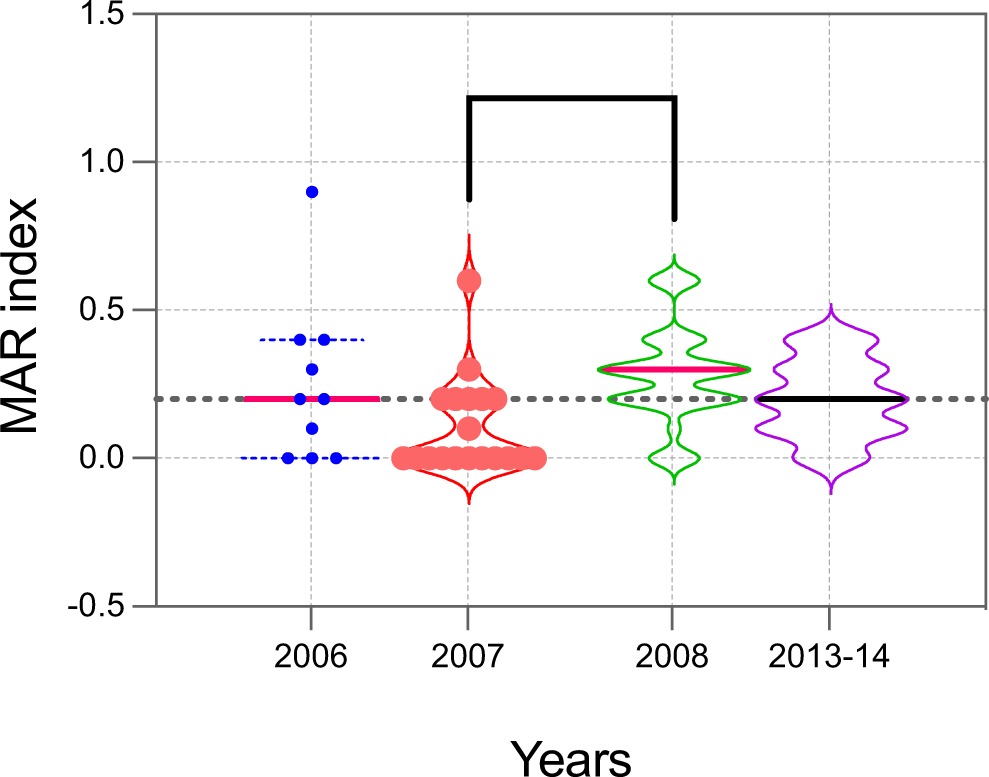
Comparison of the MAR index per year. Isolates are presented as individual values with each group’s mean and standard error plotted. MAR indicates > 0.2 (dotted line) that isolates likely originated from areas of high antibiotic use or high-risk source (90). Ordinary ANOVA was performed to compare the mean between groups. ***, P < 0.001.

## References

1. Qadri F, Svennerholm A-M, Faruque ASG, Sack RB. 2005. Enterotoxigenic Escherichia coli in developing countries: epidemiology, microbiology, clinical features, treatment, and prevention. Clinical microbiology reviews 18:465–483.

2. Kotloff KL, Nataro JP, Blackwelder WC, Nasrin D, Farag TH, Panchalingam S, Wu Y, Sow SO, Sur D, Breiman RF, Faruque AS, Zaidi AK, Saha D, Alonso PL, Tamboura B, Sanogo D, Onwuchekwa U, Manna B, Ramamurthy T, Kanungo S, Ochieng JB, Omore R, Oundo JO, Hossain A, Das SK, Ahmed S, Qureshi S, Quadri F, Adegbola RA, Antonio M, Hossain MJ, Akinsola A, Mandomando I, Nhampossa T, Acácio S, Biswas K, O’Reilly CE, Mintz ED, Berkeley LY, Muhsen K, Sommerfelt H, Robins-Browne RM, Levine MM. 2013. Burden and aetiology of diarrhoeal disease in infants and young children in developing countries (the Global Enteric Multicenter Study, GEMS): a prospective, case-control study. Lancet 382:209–22.

3. Khalil IA, Troeger C, Blacker BF, Rao PC, Brown A, Atherly DE, Brewer TG, Engmann CM, Houpt ER, Kang G, Kotloff KL, Levine MM, Luby SP, MacLennan CA, Pan WK, Pavlinac PB, Platts-Mills JA, Qadri F, Riddle MS, Ryan ET, Shoultz DA, Steele AD, Walson JL, Sanders JW, Mokdad AH, Murray CJL, Hay SI, Reiner RC, Jr. 2018. Morbidity and mortality due to shigella and enterotoxigenic Escherichia coli diarrhoea: the Global Burden of Disease Study 1990-2016. Lancet Infect Dis 18:1229–1240.

4. Del Canto F, Botkin DJ, Valenzuela P, Popov V, Ruiz-Perez F, Nataro JP, Levine MM, Stine OC, Pop M, Torres AG, Vidal R. 2012. Identification of Coli Surface Antigen 23, a novel adhesin of enterotoxigenic Escherichia coli. Infect Immun 80:2791–801.

5. Hazen TH, Nagaraj S, Sen S, Permala-Booth J, Canto FD, Vidal R, Barry EM, Bitoun JP, Chen WH, Tennant SM, Rasko DA, Overall CM. 2019. Genome and Functional Characterization of Colonization Factor Antigen I- and CS6-Encoding Heat-Stable Enterotoxin-Only Enterotoxigenic Escherichia coli Reveals Lineage and Geographic Variation. mSystems 4:e00329–18.

6. Pichel M, Binsztein N, Viboud G. 2000. CS22, a novel human enterotoxigenic Escherichia coli adhesin, is related to CS15. Infect Immun 68:3280–5.

7. Svennerholm AM, Lundgren A. 2012. Recent progress toward an enterotoxigenic *Escherichia coli* vaccine. Expert Rev Vaccines 11:495–507.

8. Isidean SD, Riddle MS, Savarino SJ, Porter CK. 2011. A systematic review of ETEC epidemiology focusing on colonization factor and toxin expression. Vaccine 29:6167–78.

9. Gonzales L, Ali ZB, Nygren E, Wang Z, Karlsson S, Zhu B, Quiding-Jarbrink M, Sjoling A. 2013. Alkaline pH Is a signal for optimal production and secretion of the heat labile toxin, LT in enterotoxigenic Escherichia coli (ETEC). PLoS One 8:e74069.

10. Gonzales L, Joffre E, Rivera R, Sjöling Å, Svennerholm AM, Iñiguez V. 2013. Prevalence, seasonality and severity of disease caused by pathogenic Escherichia coli in children with diarrhoea in Bolivia. J Med Microbiol 62:1697–1706.

11. Guzman-Otazo J, Gonzales-Siles L, Poma V, Bengtsson-Palme J, Thorell K, Flach C- F, Iñiguez V, Sjöling Å. 2019. Diarrheal bacterial pathogens and multi-resistant enterobacteria in the Choqueyapu River in La Paz, Bolivia. PLOS ONE 14:e0210735.

12. Poma V, Mamani N, Iñiguez V. 2016. Impact of urban contamination of the La Paz River basin on thermotolerant coliform density and occurrence of multiple antibiotic resistant enteric pathogens in river water, irrigated soil and fresh vegetables. SpringerPlus 5:499.

13. Begum YA, Talukder KA, Nair GB, Khan SI, Svennerholm AM, Sack RB, Qadri F. 2007. Comparison of enterotoxigenic Escherichia coli isolated from surface water and diarrhoeal stool samples in Bangladesh. Can J Microbiol 53:19–26.

14. Lothigius A, Janzon A, Begum Y, Sjöling A, Qadri F, Svennerholm AM, Bölin I. 2008. Enterotoxigenic Escherichia coli is detectable in water samples from an endemic area by real-time PCR. J Appl Microbiol 104:1128–36.

15. Wilkinson JL, Boxall ABA, Kolpin DW, Leung KMY, Lai RWS, Galbán-Malagón C, Adell AD, Mondon J, Metian M, Marchant RA, Bouzas-Monroy A, Cuni-Sanchez A, Coors A, Carriquiriborde P, Rojo M, Gordon C, Cara M, Moermond M, Luarte T, Petrosyan V, Perikhanyan Y, Mahon CS, McGurk CJ, Hofmann T, Kormoker T, Iniguez V, Guzman-Otazo J, Tavares JL, Gildasio De Figueiredo F, Razzolini MTP, Dougnon V, Gbaguidi G, Traoré O, Blais JM, Kimpe LE, Wong M, Wong D, Ntchantcho R, Pizarro J, Ying G- G, Chen C-E, Páez M, Martínez-Lara J, Otamonga J-P, Poté J, Ifo SA, Wilson P, Echeverría-Sáenz S, Udikovic-Kolic N, Milakovic M, et al. 2022. Pharmaceutical pollution of the world&#x2019;s rivers. Proceedings of the National Academy of Sciences 119:e2113947119.

16. Sjöling Å, von Mentzer A, Svennerholm AM. 2015. Implications of enterotoxigenic Escherichia coli genomics for vaccine development. Expert Rev Vaccines 14:551–60.

17. Vidal RM, Muhsen K, Tennant SM, Svennerholm AM, Sow SO, Sur D, Zaidi AKM, Faruque ASG, Saha D, Adegbola R, Hossain MJ, Alonso PL, Breiman RF, Bassat Q, Tamboura B, Sanogo D, Onwuchekwa U, Manna B, Ramamurthy T, Kanungo S, Ahmed S, Qureshi S, Quadri F, Hossain A, Das SK, Antonio M, Mandomando I, Nhampossa T, Acácio S, Omore R, Ochieng JB, Oundo JO, Mintz ED, O’Reilly CE, Berkeley LY, Livio S, Panchalingam S, Nasrin D, Farag TH, Wu Y, Sommerfelt H, Robins-Browne RM, Del Canto F, Hazen TH, Rasko DA, Kotloff KL, Nataro JP, Levine MM. 2019. Colonization factors among enterotoxigenic *Escherichia coli i*solates from children with moderate-to-severe diarrhea and from matched controls in the Global Enteric Multicenter Study (GEMS). PLoS Negl Trop Dis 13:e0007037.

18. von Mentzer A, Connor TR, Wieler LH, Semmler T, Iguchi A, Thomson NR, Rasko DA, Joffre E, Corander J, Pickard D, Wiklund G, Svennerholm AM, Sjoling A, Dougan G. 2014. Identification of enterotoxigenic Escherichia coli (ETEC) clades with long-term global distribution. Nat Genet 46:1321–6.

19. Joffré E, Iñiguez Rojas V. 2020. Molecular Epidemiology of Enteroaggregative *Escherichia coli* (EAEC) Isolates of Hospitalized Children from Bolivia Reveal High Heterogeneity and Multidrug-Resistance. Int J Mol Sci 21.

20. Zhang AN, Li L-G, Ma L, Gillings MR, Tiedje JM, Zhang T. 2018. Conserved phylogenetic distribution and limited antibiotic resistance of class 1 integrons revealed by assessing the bacterial genome and plasmid collection. Microbiome 6:130.

21. von Mentzer A, Blackwell GA, Pickard D, Boinett CJ, Joffré E, Page AJ, Svennerholm AM, Dougan G, Sjöling Å. 2021. Long-read-sequenced reference genomes of the seven major lineages of enterotoxigenic Escherichia coli (ETEC) circulating in modern time. Sci Rep 11:9256.

22. Kostakioti M, Hadjifrangiskou M, Hultgren SJ. 2013. Bacterial biofilms: development, dispersal, and therapeutic strategies in the dawn of the postantibiotic era. Cold Spring Harb Perspect Med 3:a010306.

23. Kreisberg RB, Harper J, Strauman MC, Marohn M, Clements JD, Nataro JP. 2011. Induction of increased permeability of polarized enterocyte monolayers by enterotoxigenic Escherichia coli heat-labile enterotoxin. Am J Trop Med Hyg 84:451–5.

24. Gonzales L, Sanchez S, Zambrana S, Iñiguez V, Wiklund G, Svennerholm AM, Sjöling A. 2013. Molecular characterization of enterotoxigenic Escherichia coli isolates recovered from children with diarrhea during a 4-year period (2007 to 2010) in Bolivia. J Clin Microbiol 51:1219–25.

25. Joffré E, von Mentzer A, Svennerholm A-M, Sjöling Å. 2016. Identification of new heat-stable (STa) enterotoxin allele variants produced by human enterotoxigenic Escherichia coli (ETEC). International Journal of Medical Microbiology 306:586–594.

26. Joffré E, von Mentzer A, Abd El Ghany M, Oezguen N, Savidge T, Dougan G, Svennerholm A-M, Sjöling Å. 2015. Allele variants of enterotoxigenic Escherichia coli heat-labile toxin are globally transmitted and associated with colonization factors. Journal of bacteriology 197:392–403.

27. Joffré E, Sjöling Å. 2016. The LT1 and LT2 variants of the enterotoxigenic Escherichia coli (ETEC) heat-labile toxin (LT) are associated with major ETEC lineages. Gut Microbes 7:75–81.

28. Iskandar K, Molinier L, Hallit S, Sartelli M, Hardcastle TC, Haque M, Lugova H, Dhingra S, Sharma P, Islam S, Mohammed I, Naina Mohamed I, Hanna PA, Hajj SE, Jamaluddin NAH, Salameh P, Roques C. 2021. Surveillance of antimicrobial resistance in low- and middle-income countries: a scattered picture. Antimicrobial Resistance & Infection Control 10:63.

29. Rodas C, Mamani R, Blanco J, Blanco JE, Wiklund G, Svennerholm A-M, Sjöling A, Iniguez V. 2011. Enterotoxins, colonization factors, serotypes and antimicrobial resistance of enterotoxigenic Escherichia coli (ETEC) strains isolated from hospitalized children with diarrhea in Bolivia. Braz J Infect Dis 15:132–137.

30. Gonzales-Siles L, Sjöling Å. 2016. The different ecological niches of enterotoxigenic Escherichia coli. Environmental Microbiology 18:741–751.

31. Ohno A, Marui A, Castro ES, Reyes AAB, Elio-Calvo D, Kasitani H, Ishii Y, Yamaguchi K. 1997. Enteropathogenic Bacteria in the La Paz River of Bolivia. The American Journal of Tropical Medicine and Hygiene 57:438–444.

32. Ahmed D, Islam MS, Begum YA, Janzon A, Qadri F, Sjöling Å. 2013. Presence of enterotoxigenic Escherichia coli in biofilms formed in water containers in poor households coincides with epidemic seasons in Dhaka. Journal of Applied Microbiology 114:1223–1229.

33. Brennan FP, Abram F, Chinalia FA, Richards KG, O’Flaherty V. 2010. Characterization of environmentally persistent Escherichia coli isolates leached from an Irish soil. Appl Environ Microbiol 76:2175–80.

34. Byappanahalli M, Fowler M, Shively D, Whitman R. 2003. Ubiquity and persistence of Escherichia coli in a Midwestern coastal stream. Appl Environ Microbiol 69:4549–55.

35. Maldonado Y, Fiser JC, Nakatsu CH, Bhunia AK. 2005. Cytotoxicity potential and genotypic characterization of Escherichia coli isolates from environmental and food sources. Appl Environ Microbiol 71:1890–8.

36. Power ML, Littlefield-Wyer J, Gordon DM, Veal DA, Slade MB. 2005. Phenotypic and genotypic characterization of encapsulated Escherichia coli isolated from blooms in two Australian lakes. Environ Microbiol 7:631–40.

37. Lothigius A, Sjöling A, Svennerholm AM, Bölin I. 2010. Survival and gene expression of enterotoxigenic Escherichia coli during long-term incubation in sea water and freshwater. J Appl Microbiol 108:1441–9.

38. Medina C, Ginn O, Brown J, Soria F, Garvizu C, Salazar Á, Tancara A, Herrera J. 2021. Detection and assessment of the antibiotic resistance of Enterobacteriaceae recovered from bioaerosols in the Choqueyapu River area, La Paz – Bolivia. Science of The Total Environment 760:143340.

39. Salazar D, Ginn O, Brown J, Soria F, Garvizu C. 2020. Assessment of antibiotic resistant coliforms from bioaerosol samples collected above a sewage-polluted river in La Paz, Bolivia. Int J Hyg Environ Health 228:113494.

40. Rodas C, Klena JD, Nicklasson M, Iniguez V, Sjöling A. 2011. Clonal relatedness of enterotoxigenic Escherichia coli (ETEC) strains expressing LT and CS17 isolated from children with diarrhoea in La Paz, Bolivia. PLoS One 6:e18313.

41. Gaastra W, Svennerholm AM. 1996. Colonization factors of human enterotoxigenic Escherichia coli (ETEC). Trends Microbiol 4:444–52.

42. Qadri F, Svennerholm AM, Faruque AS, Sack RB. 2005. Enterotoxigenic Escherichia coli in developing countries: epidemiology, microbiology, clinical features, treatment, and prevention. Clin Microbiol Rev 18:465–83.

43. Rasko DA, Del Canto F, Luo Q, Fleckenstein JM, Vidal R, Hazen TH. 2019. Comparative genomic analysis and molecular examination of the diversity of enterotoxigenic Escherichia coli isolates from Chile. PLOS Neglected Tropical Diseases 13:e0007828.

44. Ministerio de Medio Ambiente y Agua MdDRyT, Gobierno autonomo departamental de La Paz, Gobiernos autonomos departamentales de La Paz y Mecapaca, Autoridad de Fiscalizacion y control social de agua potable y saneamiento basico. 2017. Seguimiento a las recomendaciones del Informe de Auditoria Ambiental K2/AP05/G12 sobre los impactos negativos en la cuencia del Rio de La Paz.123.

45. Sahl JW, Sistrunk JR, Baby NI, Begum Y, Luo Q, Sheikh A, Qadri F, Fleckenstein JM, Rasko DA. 2017. Insights into enterotoxigenic Escherichia coli diversity in Bangladesh utilizing genomic epidemiology. Sci Rep 7:3402.

46. Yang C, Li Y, Zuo L, Jiang M, Zhang X, Xie L, Luo M, She Y, Wang L, Jiang Y, Wu S, Cai R, Shi X, Cui Y, Wan C, Hu Q. 2021. Genomic Epidemiology and Antimicrobial Susceptibility Profile of Enterotoxigenic Escherichia coli From Outpatients With Diarrhea in Shenzhen, China, 2015–2020. Frontiers in Microbiology 12.

47. Steinsland H, Lacher DW, Sommerfelt H, Whittam TS. 2010. Ancestral lineages of human enterotoxigenic *Escherichia coli*. J Clin Microbiol 48:2916–24.

48. Touchon M, Perrin A, de Sousa JAM, Vangchhia B, Burn S, O’Brien CL, Denamur E, Gordon D, Rocha EPC. 2020. Phylogenetic background and habitat drive the genetic diversification of Escherichia coli. PLOS Genetics 16:e1008866.

49. Naimi TS, Wicklund JH, Olsen SJ, Krause G, Wells JG, Bartkus JM, Boxrud DJ, Sullivan M, Kassenborg H, Besser JM, Mintz ED, Osterholm MT, Hedberg CW. 2003. Concurrent Outbreaks of Shigella sonnei and Enterotoxigenic Escherichia coli Infections Associated with Parsley: Implications for Surveillance and Control of Foodborne Illness†. Journal of Food Protection 66:535–541.

50. Ethelberg S, Lisby M, Böttiger B, Schultz AC, Villif A, Jensen T, Olsen KE, Scheutz F, Kjelsø C, Muller L. 2010. Outbreaks of gastroenteritis linked to lettuce, Denmark, January 2010. Eurosurveillance 15:19484.

51. Shin J, Yoon K-B, Jeon D-Y, Oh S-S, Oh K-H, Chung GT, Kim SW, Cho S-H. 2016. Consecutive Outbreaks of Enterotoxigenic Escherichia coli O6 in Schools in South Korea Caused by Contamination of Fermented Vegetable Kimchi. Foodborne Pathogens and Disease 13:535–543.

52. Shaw RK, Berger CN, Pallen MJ, Sjöling Å, Frankel G. 2011. Flagella mediate attachment of enterotoxigenic Escherichia coli to fresh salad leaves. Environmental Microbiology Reports 3:112–117.

53. Jang J, Hur HG, Sadowsky MJ, Byappanahalli MN, Yan T, Ishii S. 2017. Environmental Escherichia coli: ecology and public health implications-a review. J Appl Microbiol 123:570–581.

54. Chadha J. 2021. In vitro effects of sub-inhibitory concentrations of amoxicillin on physiological responses and virulence determinants in a commensal strain of Escherichia coli. Journal of Applied Microbiology 131:682–694.

55. Ojima Y, Nunogami S, Taya M. 2016. Antibiofilm effect of warfarin on biofilm formation of Escherichia coli promoted by antimicrobial treatment. J Glob Antimicrob Resist 7:102–105.

56. Ishii S, Sadowsky MJ. 2008. Escherichia coli in the Environment: Implications for Water Quality and Human Health. Microbes Environ 23:101–8.

57. Schaufler K, Semmler T, Wieler LH, Wöhrmann M, Baddam R, Ahmed N, Müller K, Kola A, Fruth A, Ewers C, Guenther S. 2016. Clonal spread and interspecies transmission of clinically relevant ESBL-producing Escherichia coli of ST410--another successful pandemic clone? FEMS Microbiol Ecol 92.

58. Falgenhauer L, Imirzalioglu C, Ghosh H, Gwozdzinski K, Schmiedel J, Gentil K, Bauerfeind R, Kämpfer P, Seifert H, Michael GB, Schwarz S, Pfeifer Y, Werner G, Pietsch M, Roesler U, Guerra B, Fischer J, Sharp H, Käsbohrer A, Goesmann A, Hille K, Kreienbrock L, Chakraborty T. 2016. Circulation of clonal populations of fluoroquinolone-resistant CTX-M-15-producing Escherichia coli ST410 in humans and animals in Germany. Int J Antimicrob Agents 47:457–65.

59. Ortega-Paredes D, Barba P, Mena-López S, Espinel N, Zurita J. 2018. Escherichia coli hyperepidemic clone ST410-A harboring blaCTX-M-15 isolated from fresh vegetables in a municipal market in Quito-Ecuador. International Journal of Food Microbiology 280:41–45.

60. Magaña-Lizárraga JA, Gómez-Gil B, Rendón-Maldonado JG, Delgado-Vargas F, Vega-López IF, Báez-Flores ME. 2022. Genomic Profiling of Antibiotic-Resistant Escherichia coli Isolates from Surface Water of Agricultural Drainage in North-Western Mexico: Detection of the International High-Risk Lineages ST410 and ST617. Microorganisms 10.

61. Guzman-Otazo J, Joffré E, Agramont J, Mamani N, Jutkina J, Boulund F, Hu YOO, Jumilla-Lorenz D, Farewell A, Larsson DGJ, Flach CF, Iñiguez V, Sjöling Å. 2022. Conjugative transfer of multi-drug resistance IncN plasmids from environmental waterborne bacteria to Escherichia coli. Front Microbiol 13:997849.

62. Nesse LL, Sekse C, Berg K, Johannesen KCS, Solheim H, Vestby LK, Urdahl AM. 2014. Potentially Pathogenic Escherichia coli Can Form a Biofilm under Conditions Relevant to the Food Production Chain. Applied and Environmental Microbiology 80:2042–2049.

63. Cruz-Córdova A, Espinosa-Mazariego K, Ochoa SA, Saldaña Z, Rodea GE, Cázares-Domínguez V, Rodríguez-Ramírez V, Eslava-Campos CA, Navarro-Ocaña A, Arrellano-Galindo J, Hernández-Castro R, Gómez-Duarte OG, Qadri F, Xicohtencatl-Cortes J. 2014. CS21 positive multidrug-resistant ETEC clinical isolates from children with diarrhea are associated with self-aggregation, and adherence. Frontiers in Microbiology 5.

64. Zeighami H, Haghi F, Hajiahmadi F, Kashefiyeh M, Memariani M. 2015. Multi-drug-resistant enterotoxigenic and enterohemorrhagic Escherichia coli isolated from children with diarrhea. Journal of Chemotherapy 27:152–155.

65. Belete MA, Demlie TB, Chekole WS, Sisay Tessema T. 2022. Molecular identification of diarrheagenic Escherichia coli pathotypes and their antibiotic resistance patterns among diarrheic children and in contact calves in Bahir Dar city, Northwest Ethiopia. PLOS ONE 17:e0275229.

66. Begum YA, Talukder KA, Azmi IJ, Shahnaij M, Sheikh A, Sharmin S, Svennerholm AM, Qadri F. 2016. Resistance Pattern and Molecular Characterization of Enterotoxigenic Escherichia coli (ETEC) Strains Isolated in Bangladesh. PLoS One 11:e0157415.

67. Guiral E, Gonçalves Quiles M, Muñoz L, Moreno-Morales J, Alejo-Cancho I, Salvador P, Alvarez-Martinez MJ, Marco F, Vila J. 2019. Emergence of Resistance to Quinolones and β-Lactam Antibiotics in Enteroaggregative and Enterotoxigenic Escherichia coli Causing Traveler’s Diarrhea. Antimicrob Agents Chemother 63.

68. Sati HF, Bruinsma N, Galas M, Hsieh J, Sanhueza A, Ramon Pardo P, Espinal MA. 2019. Characterizing Shigella species distribution and antimicrobial susceptibility to ciprofloxacin and nalidixic acid in Latin America between 2000-2015. PLoS One 14:e0220445.

69. Ahmed S, Korpe P, Ahmed T, Chisti MJ, Faruque ASG. 2018. Burden and Risk Factors of Antimicrobial Use in Children Less Than 5 Years of Age with Diarrheal Illness in Rural Bangladesh. Am J Trop Med Hyg 98:1571–1576.

70. Collaborators AR. 2022. Global burden of bacterial antimicrobial resistance in 2019: a systematic analysis. Lancet 399:629–655.

71. Kwak YK, Colque P, Byfors S, Giske CG, Möllby R, Kühn I. 2015. Surveillance of antimicrobial resistance among Escherichia coli in wastewater in Stockholm during 1 year: does it reflect the resistance trends in the society? Int J Antimicrob Agents 45:25–32.

72. Paulshus E, Kühn I, Möllby R, Colque P, O’Sullivan K, Midtvedt T, Lingaas E, Holmstad R, Sørum H. 2019. Diversity and antibiotic resistance among Escherichia coli populations in hospital and community wastewater compared to wastewater at the receiving urban treatment plant. Water Res 161:232–241.

73. Paulshus E, Thorell K, Guzman-Otazo J, Joffre E, Colque P, Kühn I, Möllby R, Sørum H, Sjöling Å. 2019. Repeated Isolation of Extended-Spectrum-β-Lactamase-Positive Escherichia coli Sequence Types 648 and 131 from Community Wastewater Indicates that Sewage Systems Are Important Sources of Emerging Clones of Antibiotic-Resistant Bacteria. Antimicrob Agents Chemother 63.

74. Kühn I, Möllby R. 1993. The PhP RS system. A simple microplate method for studying coliform bacterial populations. Journal of Microbiological Methods 17:255–259.

75. Blanch AR, Belanche-Muñoz L, Bonjoch X, Ebdon J, Gantzer C, Lucena F, Ottoson J, Kourtis C, Iversen A, Kühn I, Mocé L, Muniesa M, Schwartzbrod J, Skraber S, Papageorgiou GT, Taylor H, Wallis J, Jofre J. 2006. Integrated analysis of established and novel microbial and chemical methods for microbial source tracking. Appl Environ Microbiol 72:5915–26.

76. Page AJ, De Silva N, Hunt M, Quail MA, Parkhill J, Harris SR, Otto TD, Keane JA. 2016. Robust high-throughput prokaryote de novo assembly and improvement pipeline for Illumina data. Microbial Genomics 2.

77. Zerbino DR, Birney E. 2008. Velvet: Algorithms for de novo short read assembly using de Bruijn graphs. Genome Research 18:821–829.

78. Boetzer M, Henkel CV, Jansen HJ, Butler D, Pirovano W. 2010. Scaffolding pre-assembled contigs using SSPACE. Bioinformatics 27:578–579.

79. Nadalin F, Vezzi F, Policriti A. 2012. GapFiller: a de novo assembly approach to fill the gap within paired reads. BMC Bioinformatics 13:S8.

80. Seemann T. 2014. Prokka: rapid prokaryotic genome annotation. Bioinformatics 30:2068–9.

81. Pruitt KD, Tatusova T, Brown GR, Maglott DR. 2011. NCBI Reference Sequences (RefSeq): current status, new features and genome annotation policy. Nucleic Acids Research 40:D130–D135.

82. Beghain J, Bridier-Nahmias A, Le Nagard H, Denamur E, Clermont O. 2018. ClermonTyping: an easy-to-use and accurate in silico method for Escherichia genus strain phylotyping. Microb Genom 4.

83. Hunt M, Mather AE, Sánchez-Busó L, Page AJ, Parkhill J, Keane JA, Harris SR. 2017. ARIBA: rapid antimicrobial resistance genotyping directly from sequencing reads. Microb Genom 3:e000131.

84. Page AJ, Cummins CA, Hunt M, Wong VK, Reuter S, Holden MT, Fookes M, Falush D, Keane JA, Parkhill J. 2015. Roary: rapid large-scale prokaryote pan genome analysis. Bioinformatics 31:3691–3.

85. Page AJ, Taylor B, Delaney AJ, Soares J, Seemann T, Keane JA, Harris SR. 2016. SNP-sites: rapid efficient extraction of SNPs from multi-FASTA alignments. Microb Genom 2:e000056.

86. Yu G. 2020. Using ggtree to Visualize Data on Tree-Like Structures. Current Protocols in Bioinformatics 69:e96.

87. Katoh K, Misawa K, Kuma Ki, Miyata T. 2002. MAFFT: a novel method for rapid multiple sequence alignment based on fast Fourier transform. Nucleic Acids Research 30:3059–3066.

88. Colque Navarro P, Fernandez H, Möllby R, Otth L, Tiodolf M, Wilson M, Kühn I. 2014. Antibiotic resistance in environmental Escherichia coli – a simple screening method for simultaneous typing and resistance determination. Journal of Water and Health 12:692–701.

89. Krumperman PH. 1983. Multiple antibiotic resistance indexing of Escherichia coli to identify high-risk sources of fecal contamination of foods. Appl Environ Microbiol 46:165–70.

90. Paul S, Bezbaruah RL, Roy MK, Ghosh AC. 1997. Multiple antibiotic resistance (MAR) index and its reversion in Pseudomonas aeruginosa. Lett Appl Microbiol 24:169–71.

91. Joffre E, Nicklasson M, Álvarez-Carretero S, Xiao X, Sun L, Nookaew I, Zhu B, Sjöling Å. 2019. The bile salt glycocholate induces global changes in gene and protein expression and activates virulence in enterotoxigenic Escherichia coli. Scientific Reports 9:108.

